# Western diet increases COVID-19 disease severity in the Syrian hamster

**DOI:** 10.1101/2021.06.17.448814

**Authors:** Julia R. Port, Danielle R. Adney, Benjamin Schwarz, Jonathan E. Schulz, Daniel E. Sturdevant, Brian J. Smith, Victoria A. Avanzato, Myndi G. Holbrook, Jyothi N. Purushotham, Kaitlin A. Stromberg, Ian Leighton, Catharine M. Bosio, Carl Shaia, Vincent J. Munster

**Affiliations:** Laboratory of Virology, National Institute of Allergy and Infectious Diseases, National Institutes of Health, Hamilton, MT, USA; Laboratory of Bacteriology, National Institute of Allergy and Infectious Diseases, National Institutes of Health, Hamilton, MT, USA; Genomics Unit, Research Technologies Branch, National Institute of Allergy and Infectious Diseases, National Institutes of Health, Hamilton, MT, USA; Rocky Mountain Veterinary Branch, Division of Intramural Research, National Institutes of Health, Hamilton, MT, USA

**Keywords:** Syrian hamster, SARS-CoV-2, obesity, pathogenesis, lipid metabolism

## Abstract

Pre-existing comorbidities such as obesity or metabolic diseases can adversely affect the clinical outcome of COVID-19. Chronic metabolic disorders are globally on the rise and often a consequence of an unhealthy diet, referred to as a Western Diet. For the first time in the Syrian hamster model, we demonstrate the detrimental impact of a continuous high-fat high-sugar diet on COVID-19 outcome. We observed increased weight loss and lung pathology, such as exudate, vasculitis, hemorrhage, fibrin, and edema, delayed viral clearance and functional lung recovery, and prolonged viral shedding. This was accompanied by an increased trend of systemic IL-10 and IL-6, as well as a dysregulated serum lipid response dominated by polyunsaturated fatty acid-containing phosphatidylethanolamine, recapitulating cytokine and lipid responses associated with severe human COVID-19. Our data support the hamster model for testing restrictive or targeted diets and immunomodulatory therapies to mediate the adverse effects of metabolic disease on COVID-19.

## Introduction

Severe acute respiratory syndrome coronavirus-2 (SARS-CoV-2) is the etiological agent of coronavirus disease (COVID)-19 and can cause asymptomatic to severe lower respiratory tract infections in humans (Nie et al., 2020; Parry et al., 2020). Pre-existing comorbidities such as immunosuppression, obesity, diabetes, or chronic lung disease can adversely affect the clinical outcome (Butler and Barrientos, 2020; Hussain et al., 2020; Li et al., 2009; Petrakis et al., 2020). Of these, obesity and metabolic disorders are global pandemics of rising concern (Araújo et al., 2019; Saklayen, 2018; Swinburn et al., 2011). The underlying disease is driven mainly by changes in the global food system, which is producing more processed, affordable, and effectively marketed food than ever before. This diet, rich in saturated fats and refined sugars, is referred to as a Western Diet (Cordain et al., 2005). Long-term consumption of a Western Diet may result in chronic activation of the immune system, impairing both innate and adaptive responses (Green and Beck, 2017a, b; Rogero and Calder, 2018). The Western Diet has been associated with non-alcoholic steatohepatitis (NASH) and non-alcoholic fatty liver disease (NAFLD). These disease syndromes predispose individuals to multiple comorbidities that can include cirrhosis and liver failure. The relative risk of hospitalization and severe COVID-19 outcome are significantly increased for patients afflicted by these comorbidities (Butler and Barrientos, 2020). This has resulted in disproportionately worse outcomes in US ethnic and racial minorities, where prevalence and incidence of metabolic disorders are increased (Cefalu and Rodgers, 2021).

It is currently unclear how certain comorbidities may determine disease manifestation of COVID-19. Different studies have demonstrated that the Syrian hamster model is suitable to model aspects of obesity and diabetes and for studying lipid metabolism (Dalbøge et al., 2015; Kasim-Karakas et al., 1996). In healthy hamsters, SARS-CoV-2 infection is associated with mild to moderate clinical disease (Chan et al., 2020; Rosenke et al., 2020; Sia et al., 2020). However, no studies have investigated COVID-19 in hamsters with comorbidities. Here we show in a Syrian hamster model how a continuous high-fat high-sugar (HFHS) diet changed the metabolomic state in the Syrian hamster and the resulting consequences on viral replication dynamics, immune protection and disease severity after infection with SARS-CoV-2.

## Results

### High-fat and high-sugar diet induces metabolic changes characterized by increased early weight gain and glucose tolerance

We investigated the impact of a consistent high-fat and high-sugar (HFHS) diet on the Syrian hamster. Either a regular rodent (RD) diet or a high-calorimetric HFHS diet was given to male Syrian hamsters (4-6 week old) for 16 weeks *ad libitum* (N = 35, respectively). Weight gain of juvenile hamsters was monitored weekly. Initially, animals on the HFHS diet gained weight faster than animals on the regular diet. Difference in median weights was significant from the 2^nd^ week onwards until week 10 (Fig 1 A, N = 35, ordinary two-way ANOVA, followed by Sidak’s multiple comparisons test, p = 0.001, p = <0.001, p = <0.001, p = <0.001, p = <0.001, p = <0.001, p = <0.001, p = 0.0011, p = <0.001). After week 10 weight gain either plateaued or decreased in the HFHS group (median = 165 g), while in the regular diet group weight increased until week 12 (median = 160 g), at which point the median weight between groups showed no significant difference. We observed morbidity (4/35 = 11%) in the HFHS group, which was absent in the RD group.

**Figure 1:**
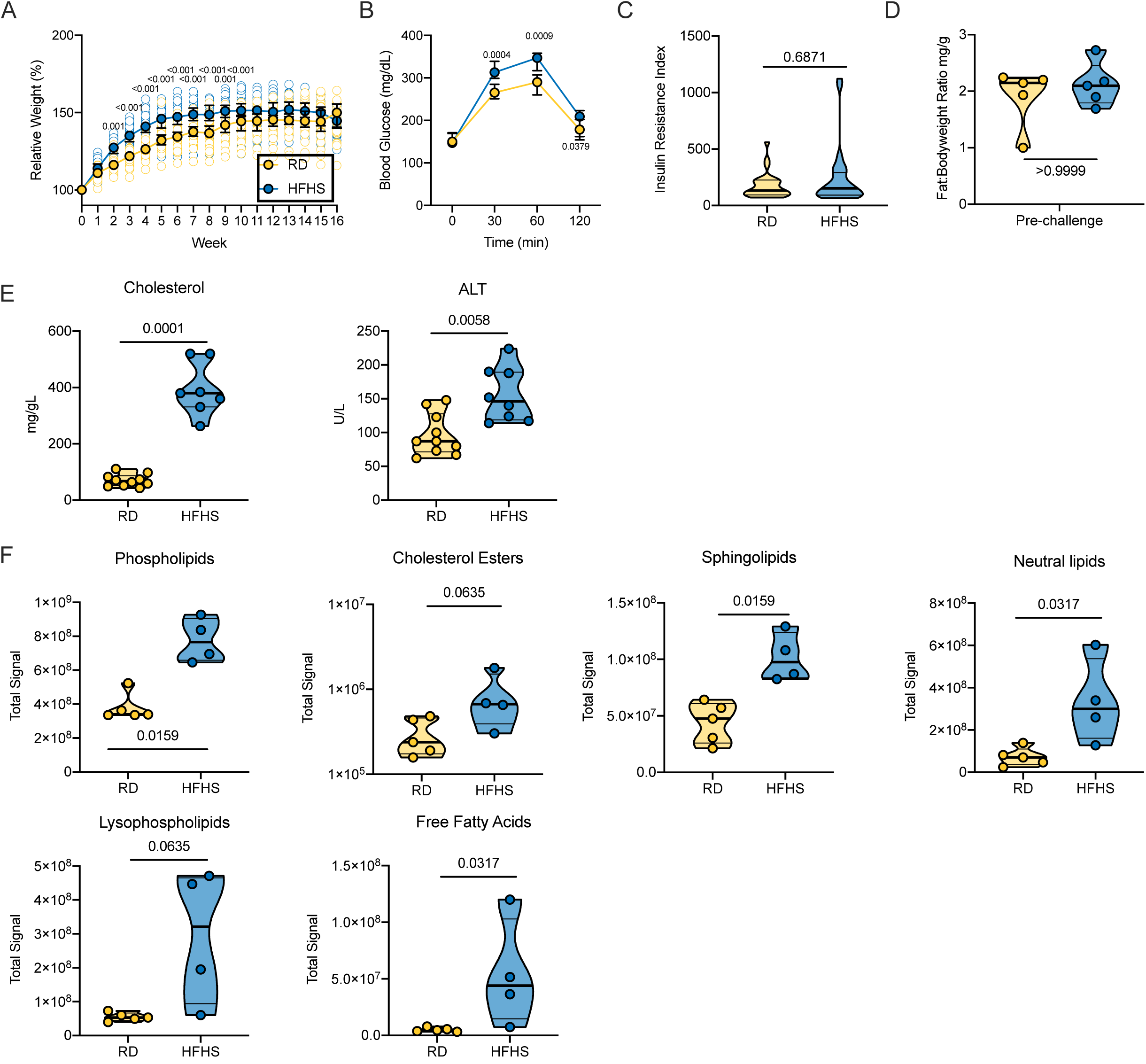
High-fat and high-sugar diet induces metabolic changes characterized by increased juvenile weight gain and glucose tolerance. Male Syrian hamsters were fed either a regular or high-fat high-sugar diet *ad libitum* for 16 weeks. **A.** Relative weight gain in hamsters on each diet regimen, measured weekly. Graphs show median ± 95% CI, N = 35, ordinary two-way ANOVA, followed by Sidak’s multiple comparisons test. **B.** Oral glucose tolerance test performed at 16 weeks. Graphs show median ± 95% CI, N = 30 (RD) / 29 (HFHS), ordinary two-way ANOVA, followed by Sidak’s multiple comparisons test. **C.** Insulin response after application of oral glucose load as shown by insulin resistance index (fasting glucose level (mmol/L) x fasting insulin level (mIU/L)). Truncated violin plots depicting median, quartiles and individuals, N = 30 (RD) / 29 (HFHS), Mann-Whitney test. **D.** Adiposity index as measured by testicular fat pads/total body weight at 16 weeks. Truncated violin plots depicting median, quartiles and individuals, N = 5, Mann-Whitney test. **E.** Blood lipid ALT and cholesterol levels measured on a commercially available lipid panel on an automated blood chemistry analyzer. **F.** Serum aggregate lipids signal analyzed by liquid chromatography tandem mass spectrometry (LC-MS/MS) at 16 weeks of diet regimen. Truncated violin plots depicting median, quartiles and individuals, N = 5(RD) / 4 (HFHS), Mann-Whitney test. Abbreviations: RD = regular diet, HFHS = high-fat high-sugar, ALT = alanine aminotransaminase. p-values are indicated were appropriate.

To assess the levels of glucose-associated symptoms triggered by a HFHS diet we conducted an oral glucose tolerance test (OGTT). No difference in fasting blood glucose levels between diet groups was observed (N = 30 (RD) / 29 (HFHS), median = 150 / 147 mg/dL). However, HFHS animals demonstrated impaired glucose intolerance upon application of an oral glucose dose; blood glucose levels 30 min, 60 min and 120 min after oral application were significantly increased compared to RD animals (Fig 1 B, N = 30 (RD) / 29 (HFHS), 30 minutes median = 265 / 313 mg/dL and 60 minutes median = 290 / 347 mg/dL, ordinary two-way ANOVA, followed by Sidak’s multiple comparisons test, p = 0.0004, p = 0.0009, respectively). We compared the insulin response after application of oral glucose load and found no difference between the diet regimens. The insulin resistance index (fasting glucose level (mmol/L) x fasting insulin level (mIU/L) showed no significant differences (Fig 1 C, N = 30 (RD) / 29 (HFHS), Mann-Whitney test, p = 0.6871) (Hayashi et al., 2013; Li et al., 2009). Five animals were euthanized pre-challenge in order to assess diet induced pathology. There was no difference in body fat-to-weight ratio (Fig 1 D, N = 5, median = 1.905 (RD) / 2.117 (HFHS) Fat:Bodyweight ratio (mg/g), Mann-Whitney test, p > 0.9999).

### High-fat and high-sugar diet induces liver damage and systemic hyperlipidemia

We investigated the changes in lipid metabolism through a blood lipid biochemistry panel (Sup Table 1). Due to increased levels of fat in the samples collected from HFHS animals, HDL and LDL could not be assessed due to incompatibility with the instrument. Total cholesterol was significantly increased in the HFHS group (Fig 1 E, N = 10 (RD) / 7 (HFHS), median = 67.6 / 380 mg/dL, Mann-Whitney test, p = 0.0001). The median (146 U/L) alanine aminotransferase (ALT), an indication of hepatocellular injury without overt cholestasis, values in the HFHS animals were above the upper limit of previously established reference ranges (Washington and Van Hoosier, 2012). To understand which lipids were circulating in serum, we analyzed serum by liquid chromatography tandem mass spectrometry (LC-MS/MS). Aggregate signals across all lipid classes assayed in the HFHS animals compared to RD were increased, comprising phospholipids, cholesterol esters, sphingolipids, neutral lipids, lysophospholipids, and free fatty acids (Fig 1 F, N = 5(RD) / 4 (HFHS), Mann-Whitney test, p = 0.0159, p = 0.0635, p = 0.0159, p = 0.0317, p = 0.0653, p = 0.0317, respectively). Hence, we further assessed changes in the liver through gross and histologic pathology. Gross pathology of livers differed substantially. Livers from animals on the HFHS diet were diffusely pale, friable, and sections floated in formalin while RD hamster livers appeared grossly normal. Histologically, hepatocytes were expanded by micro and macrovesicles in HFHS animals, while hepatocytes in RD animals appeared normal (Fig 2 A - F).

**Figure 2.**
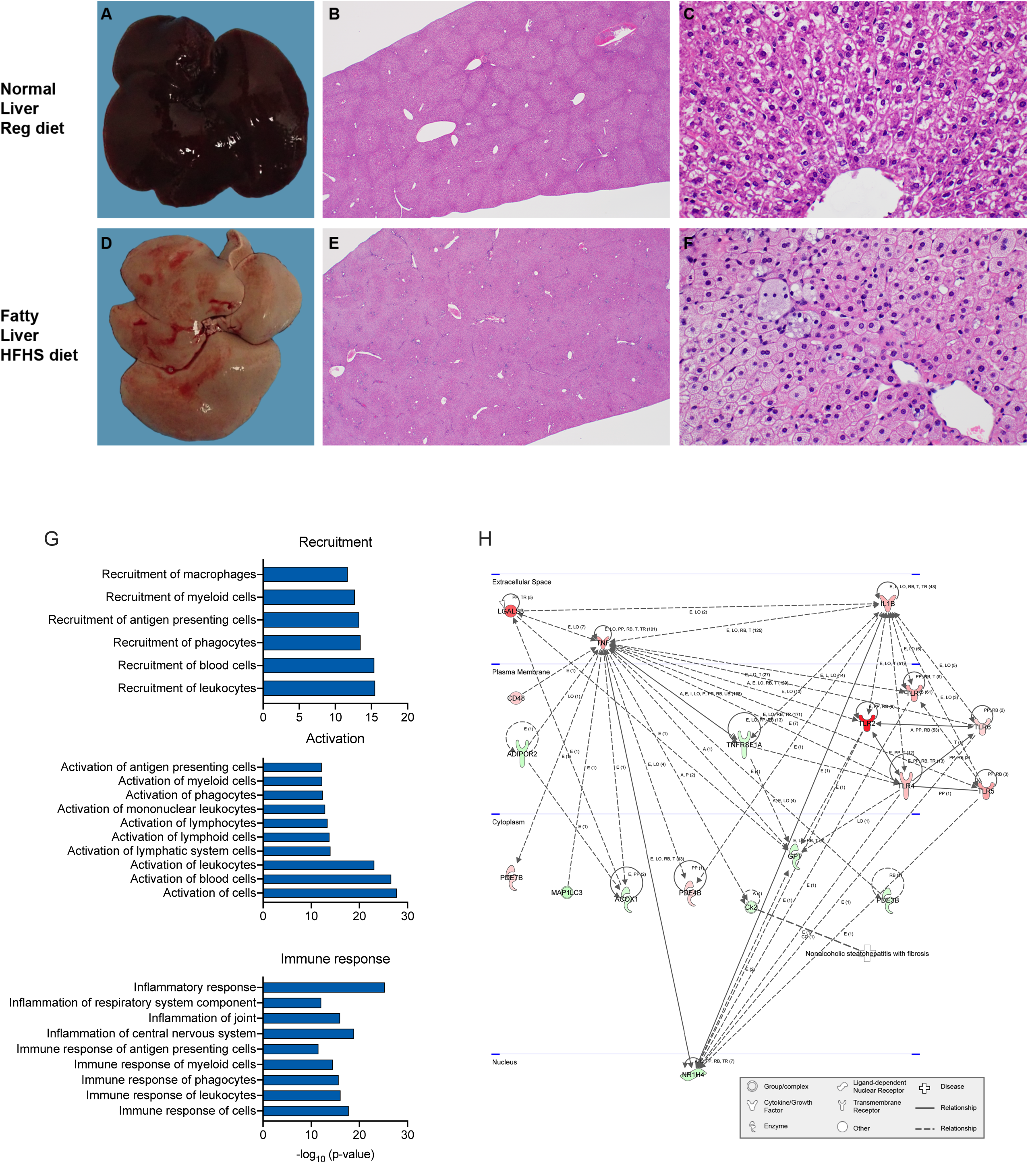
High-fat and high-sugar diet induces liver damage and systemic hyperlipidemia. Male Syrian hamsters were fed either a regular or high-fat high-sugar diet *ad libitum* and 5 animals from each group were sacrificed week 16 for analyses of liver tissue. **A.D.** Gross imaging of a representative liver from one hamster on the RD and one hamster on the HFHS diet regimen. **B.E.** 20x photomicrograph of H&E-stained slide. **C.F.** 400x photomicrograph of H&E-stained slide. **G.** RNA was isolated for gene expression analyses from liver tissue at 16 weeks. Using Integrated Pathway Analysis (Qiagen), significantly up-regulated canonical pathways were identified. Graphs show pathways associated with cell recruitment, activation, and immunological inflammation (p > 0.05, z-score < -2 or > 2). **H.** Integrated Pathway Analysis (Qiagen) was used to depict the gene network associated with nonalcoholic steatohepatitis. Symbols refer to legend below figure. Red: Gene upregulation in high-fat high-sugar animals as compared to regular diet animals. Green: downregulation in comparison to regular diet.

To further characterize the effect of the HFHS diet regimen on the liver, we evaluated global changes in the gene expression after 16 weeks. Principal components analysis of the complete gene expression profile revealed expected grouping with each diet regimen group containing their associated replicates (Sup Fig 1 A, N = 5 (RD), 4 HFHS). In total, 2,114 genes were significantly, differentially expressed (p<0.05 and >2fold) in the liver. To assess the enrichment of these differential genes, they were imported into Ingenuity Pathway Analysis (IPA) software. The results show that in the comparison of HFHS to RD animals 124 canonical pathways were significantly enriched and 200 downstream effects were predicted on biological processes and disease or toxicological function (p-value < 0.05, z-score <= -2 or >= 2): amongst which were cell recruitment, inflammation, activation, and immune-associated pathways (Fig 2 G, Sup Table 2 shows all significant predicted downstream effects). Interestingly, we also observed a pathway activation pattern reminiscent of NAFLD TNF-driven inflammation, (Fig 2 H).

Together, these data suggest that HFHS diet induced drastic changes in glucose uptake and lipid metabolism, characterized by systemic dyslipidemia and gross changes in liver pathology. This translated into increased inflammation and a gene expression profile in the liver reminiscent of fatty liver disease.

### High-fat and high-sugar diet exacerbated disease severity after SARS-CoV-2 infection

We challenged hamsters (RD: N = 20, HFHS = 13 (Group size adjusted for the HFHS group due to the morbidity of the model pre-challenge)) with 8×10^4^ TCID_50_ SARS-CoV-2 via the intranasal route. Animals were euthanized at 7 days-post inoculation (DPI) (RD: N = 10, HFHS = 4), at 14 DPI (RD: N = 5, HFHS = 4) or monitored until 21 DPI (RD: N = 5, HFHS = 5). We observed a trend of more severe morbidity in the HFHS group, in which two animals reached euthanasia criteria (>20% relative body weight loss) at 8 and 9 DPI, respectively (Fig 3 A). While the HFHS animals demonstrated non-infection associated morbidity, the timing and symptoms associated with these fatalities suggest that they were caused by the infection. In the RD group, a median peak weight loss was observed at 6 DPI (∼7% relative body weight), after which animals recovered and returned to pre-challenge weights by 14 DPI. Weight in HFHS animals was significantly decreased after 3 DPI and negative area under the curve (AUC) analysis between 1-14 DPI revealed significant difference (Fig 3 B, N = 10 (RD) / 7 (HFHS), Mann-Whitney test, p = 0.0002). In the HFHS group median peak weight loss was reached at 8 DPI (∼16% relative body weight) and no animal recovered pre-challenge weights until the end of the study at 21 DPI.

**Figure 3:**
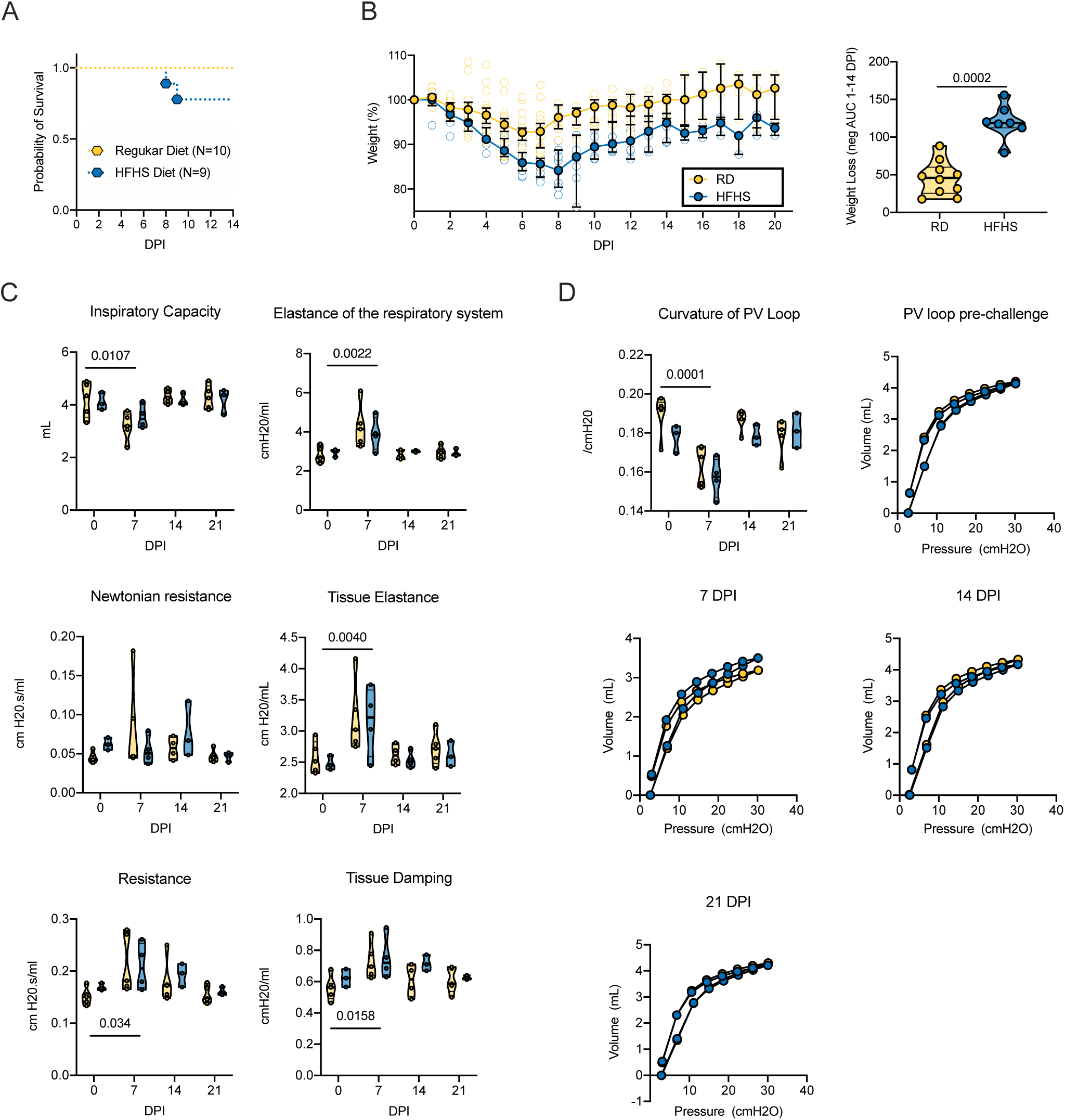
High-fat and high-sugar diet exasperated disease severity after SARS-COV-2 infection. Male Syrian hamsters were fed either a regular or high-fat high-sugar diet *ad libitum* for 16 weeks, then challenged with 8×10^4^ TCID_50_ SARS-CoV-2. **A.** Survival after challenge for RD (N = 10) and HFHS (N = 9) in the 14 and 21 DPI groups **B.** Relative weight loss in hamsters after challenge. Left graph shows median ± 95% CI. Right graph shows area under the curve (AUC, negative peaks only) between 1-14 DPI of surviving animals. Truncated violin plots depicting median, quartiles and individuals, N = 10 (RD)/ 7 (HFHS), Mann-Whitney test. **C.** Lung function analysis after challenge **D**. Pressure-volume loops at pre-challenge, 7, 14, and 21 DPI. Abbreviations: RD = regular diet, HFHS = high-fat high-sugar, DPI = days post inoculation. p-values are indicated were appropriate.

To better understand the clinical impact of a HFHS diet on SARS-CoV-2 infection, the respiratory function of the hamsters was evaluated. We performed forced oscillation tests on mechanically ventilated hamsters pre-challenge, and on 7, 14, and 21 DPI. No significant differences in pulmonary function were detected between the RD and HFHS groups at any time point.

Pulmonary function after SARS-CoV-2 infection has not been assessed in the Syrian hamster yet, so we combined the groups to evaluate changes over the course of infection. Inspiratory capacity was significantly decreased in 7 DPI as compared to pre-challenge (Figure 3 C, baseline: N = 5 (RD) / 3 (HFHS) and 7 DPI: N = 5 (RD) / 4 (HFHS), baseline median = 4.345 / 4.032 and 7 DPI median = 3.195 / 3.464 mL, ordinary two-way ANOVA, followed by Tukey’s multiple comparisons test, p = 0.0107). Elastance of the respiratory system was significantly increased at 7 DPI (baseline median = 2.68 / 3.032 and 7 DPI median = 4.138 / 3.852 cmH2O/mL, p = 0.0022), as was tissue elastance (baseline median = 2.514 / 2.450 and 7 DPI median = 3.021 / 3217 cmH2O/mL, p = 0.0040). The resistance of the airway not associated with gas exchange (Newtonian resistance) was not significantly different at any time point; however total resistance was significantly increased in 7 DPI as compared to pre-challenge (baseline median = 0.151 / 0.167 and 7 DPI median = 0.181 / 0.205 cmH2O.s/mL, p = 0.034). Changes in peripheral resistance were also detected by an increase in tissue damping at 7 DPI as compared to pre-challenge animals, which reflects how oscillatory energy is dispersed or retained within parenchymal tissue (baseline median = 0.564 / 0.623 and 7 DPI median = 0.695 / 0.720 cmH2O/mL, p = 0.0158). Recovery to pre-challenge was observed for all parameters by 14 DPI. Together, these changes in respiratory function led to an overall decrease in shape parameter k, which reflects the curvature of the pressure-volume curve, on 7 DPI (Fig 3 D, baseline median = 0.193 / 0.180 and 7 DPI median = 0.168 / 0.158 /cmH_2_0, ordinary two-way ANOVA, followed by Sidak’s multiple comparisons test, p = 0.0001). While not significant, a slower trend of recovery to pre-challenge values for resistance and tissue damping was observed in the HFHS group. This could indicate that functional lung recovery in this group was slower.

### High-fat and high-sugar diet is associated with exudate, vasculitis, inflammation of the epithelia and hemorrhage, fibrin and edema, and decreased viral clearance

Next, we assessed the pathology in the lungs at necropsy, 7 DPI. Grossly, lungs displayed lesions with multifocal dark red foci visible on the surface of the lobes (Fig 4 A-J). Across groups the 7 DPI lungs were more turgid, failed to collapse and had increased lung weights as compared to pre-challenge lungs (Sup Fig 2 A). Lung weight recovery appeared slower in HFHS animals. Histopathologically, only a subset of RD animals demonstrated increased lung damage (N = 5/10, > 50% lung tissue affected). At 7 DPI, foci were multifocal and adjacent to bronchi and blood vessels as well as peripherally along the sub pleural margin. Overall, no significant difference was seen between the cumulative pathological score between diet groups. However, three out of four animals demonstrated lesions in >50% of tissue (Fig 4 K, Sup Fig 2 B). In HFHS animals, foci were multifocal but less clearly delineated due to hemorrhage, edema, and fibrin. Interstitial pneumonia was characterized by thickened septa due to inflammatory cells, fibrin and edema and lined by hyperplastic type II pneumocytes. Alveoli were filled with inflammatory cells, edema and organizing fibrin. The two HFHS animals which were euthanized at day 8/9 due to severe disease and weight loss (>20%) both showed pneumonia, hemorrhage, edema, and inflammation (Sup Fig 3).

**Figure 4.**
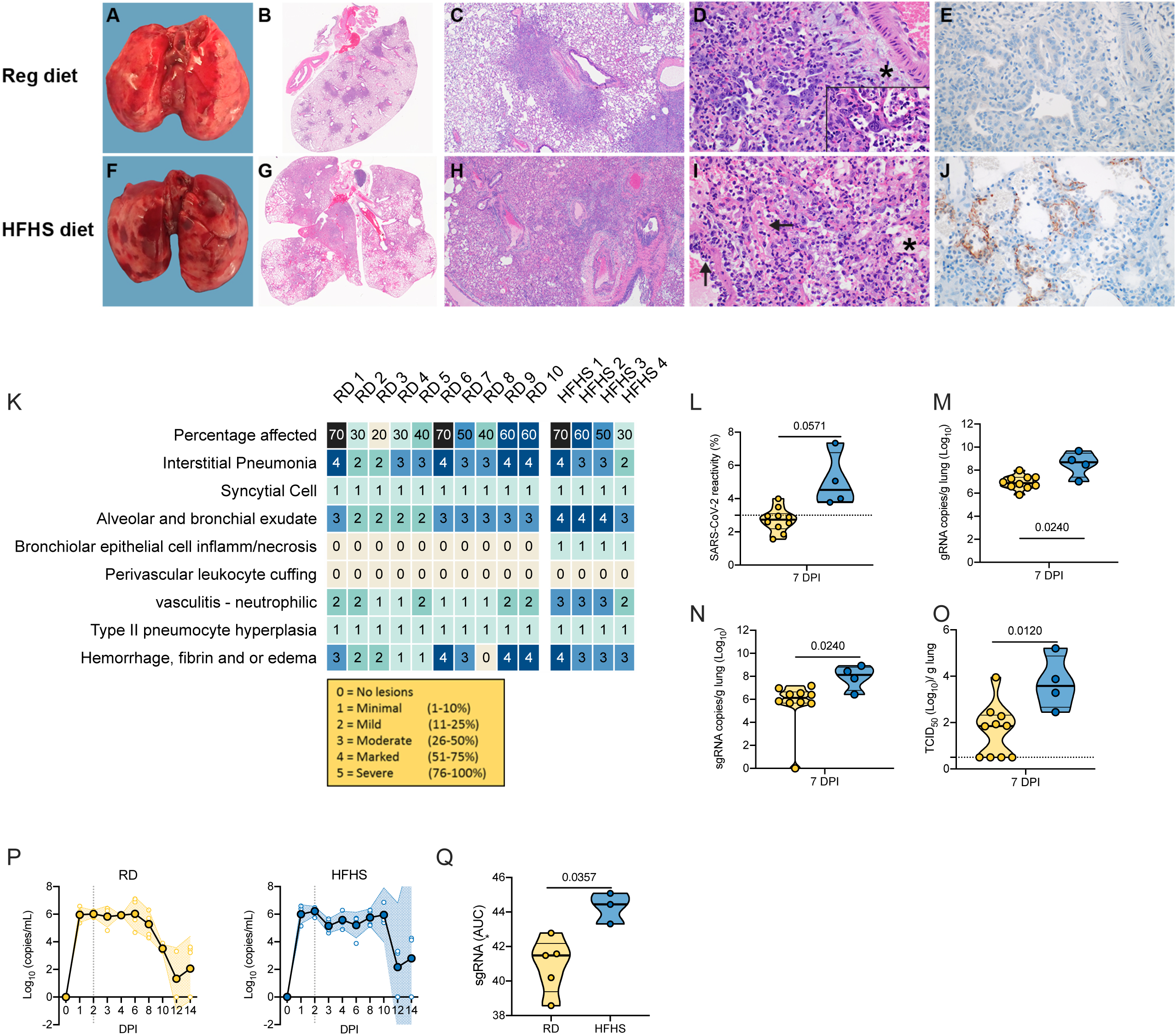
High-fat and high-sugar diet is associated to increased pulmonary pathology and decreased viral clearance. Animals were euthanized at 7 DPI with SARS-CoV-2 in order to compare lung pathology and viral titers. **A-J.** Gross and photomicrographic images of hamster lungs taken at 7 DPI. **A, F.** Gross necropsy findings consisted of multifocal well-circumscribed dark red foci throughout turgid lobes which failed to collapse. **B, G.** Dark red foci in the gross images correlate with the consolidated foci adjacent to airways and scattered along the pleural margin in the sub-gross images. HE, 1.4x. **C, H.** Foci of interstitial pneumonia adjacent to terminal bronchioles and accompanying blood vessels. HE, 20x. **D, I.** Pneumonia consists of alveoli containing neutrophils, eosinophils, alveolar and septal macrophages, fibrin, edema and septa lined by hyperplastic type II pneumocytes, HE 400x. Syncytial cells are common (see inset, HE, 1000x). Pneumonic areas in the HFHS diet hamsters frequently had abundant intra–alveolar edema (*) and organizing fibrin mixed with inflammatory cells. Note the vessel wall disrupted by sub-endothelial leukocytes and cellular debris (←). **E, J.** anti-SARS-CoV-2 immunoreactivity in the lungs from the regular diet hamsters is rare compared to the frequent pneumocyte immunoreactivity in the lungs of the HFHS diet hamsters, IHC, 400x. **K.** Individual pathological scores. **L.** Quantitative count of SARS-CoV-2 immunoreactivity by morphometric analysis. Truncated violin plots depicting median, quartiles and individuals, N = 10 (RD) / 4 (HFHS), Mann-Whitney test. **M.N.** Lung viral load measured by g and sgRNA. Truncated violin plots depicting median, quartiles and individuals, N = 10 (RD) / 4 (HFHS), Mann-Whitney test. **O.** Infectious virus measured by lung titration. Truncated violin plots depicting median, quartiles and individuals, N = 10 (RD) / 4 (HFHS), Mann-Whitney test. Dotted line = limit of detection. Abbreviations: g = genomic, sg = subgenomic, DPI = days post inoculation, H&E = hematoxylin and eosin stain, IHC = immunohistochemistry. p-values are indicated were appropriate.

At 14 DPI, thickened septa, presumably from interstitial fibrosis with alveolar bronchiolization, were observed in lungs from RD animals (N = 2) (Sup Fig 4 A-D). In contrast, HFHS animals at 14 DPI had less septal thickening and more septal, alveolar, and perivascular inflammation (N = 2). At 21 DPI four out of five of the RD animals and three out of three of the HFHS animals had thickened alveolar septa with alveolar bronchiolization (Sup Fig 4 E-H).

Immunohistochemistry staining for SARS-CoV-2 antigen was increased at 7 DPI in lungs of HFHS animals compared to RD animals (median = 2.71 (RD) / 5.043 (HFHS), N = 10 / 4) (Fig 4 E.J.L). To confirm this finding, we compared genomic RNA, subgenomic (sg)RNA (surrogate for replicating virus) and infectious viral particles isolated from lungs at 7 DPI. Levels of gRNA and sgRNA in the lungs of HFHS animals at 7 DPI were significantly increased as compared to RD animals. Additionally, no infectious virus could be isolated from a subset of RD animals and overall, significantly more infectious virus could be isolated in HFHS animals (Fig 4 M.N.O; RD: N = 10, HFHS: N = 4, gRNA median = 6.935 / 8.513 copies/g lung (log_10_), sgRNA median = 5.639 / 7.896 copies/g lung (log_10_) and infectious virus median = 1.63 / 3.703 TCID_50_/g (log_10_), Mann-Whitney test, p = 0.0240, p = 0.0240 and p = 0.0120, respectively).

To better understand if the HFHS diet contributed to changes in viral replication kinetics in the upper respiratory tract, swabs from the oropharynx were analyzed for the presence of sgRNA. Respiratory shedding in both groups peaked at 2 DPI. Shedding in HFHS animals was constantly high up until 10 DPI, while shedding began decreasing in RD animals after 6 DPI. To compare the overall shedding burden, we performed an area under the curve (AUC) analysis for both groups depicting the cumulative shedding. HFHS animals presented significantly higher cumulative shedding (Fig 4 P.Q, N = 5 (RD) / 3 (HFHS), median 41.48 / 44.44 AUC (log_10_), Mann-Whitney test, p = 0.0357).

### Immune infiltration in the lung during the acute-phase of infection and humoral immunity are not significantly affected by high-fat high-sugar diet

Using immunohistochemistry, we investigated the infiltration of macrophages (IBA 1 staining), T-cells (CD3 staining), and B-cells (Pax 5 staining) over the course of infection (Fig 5). Macrophages were detected throughout all sections but were increased in 7 and 14 DPI samples in pneumonic areas irrespective of diet regimen. In addition, T lymphocytes were increased in 7 and 14 DPI samples in pneumonic areas. No increase in B cells was observed. To quantify the influx of macrophages and T cells we used morphometric analysis (Sup Fig 5). No significant difference was seen between the RD and HFHS groups. Both macrophages and T cells increased in numbers at 7 DPI as compared to pre-challenge conditions for both groups. (Fig 6 A.B, pre-challenge: N = (RD) / 2 (HFHS) and 7 DPI: N = 10 (RD) / 4 (HFHS), median macrophages = (3.075 / 3.530 (pre-challenge)) / (13.630 / 10.480 (7 DPI)) % reactivity and median T cells = (4.515 / 4.125 (pre-challenge)) / (11.340 / 11.255 (7 DPI)) % reactivity, ordinary two-way ANOVA, followed by Sidak’s multiple comparisons test, p = 0.1007 / 0.3564 and p = 0.0001 / 0.0001, respectively).

**Figure 5.**
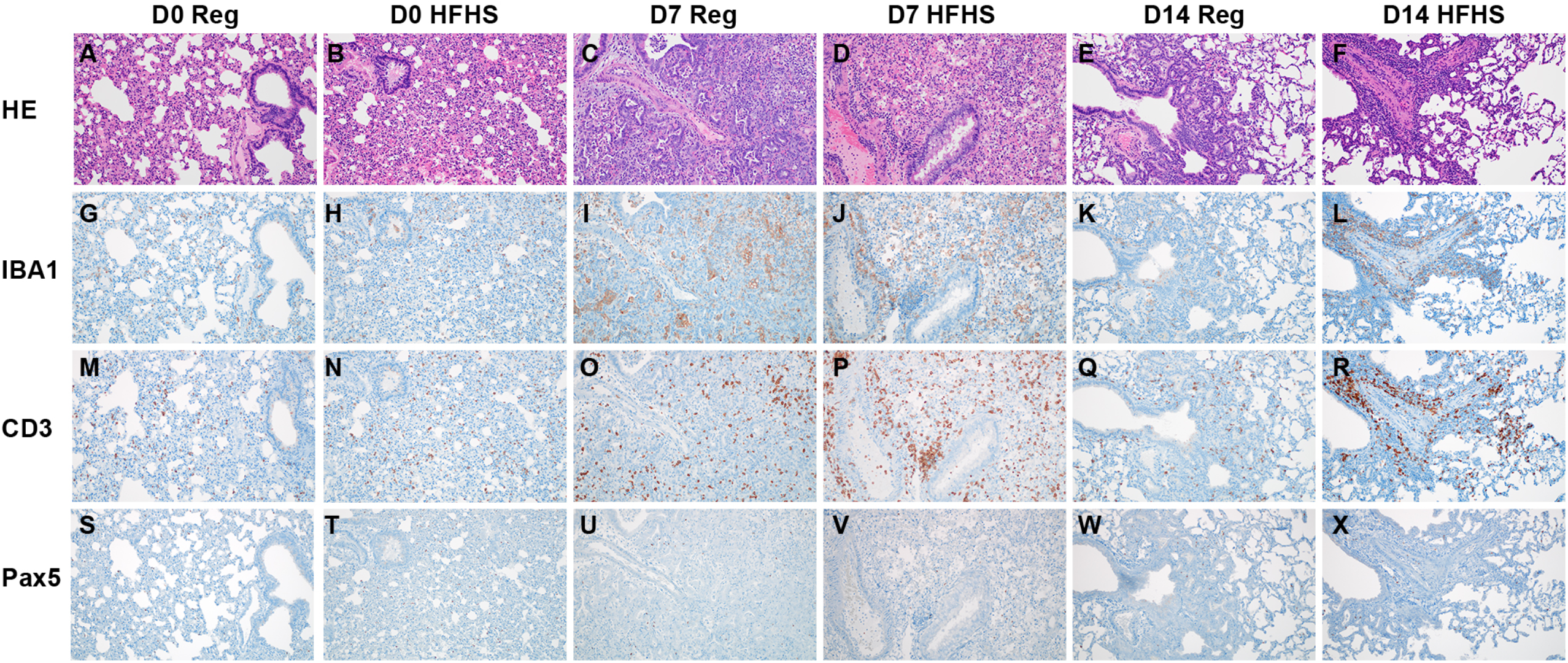
Immune infiltration and in the lung during acute-phase of infection and humoral immunity is not significantly affected by high-fat high-sugar diet. Animals were euthanized at 0, 7 and 14 DPI and the presence of SARS-CoV-2 antigen, T-cells, B-cells and macrophages investigated. **A, B.** Pre-challenge RD and HFHS diet hamster lungs. **G, H.** IBA1; **M, N.** CD3 and **S, T.** Pax5. **C, D.** Lungs at 7 DPI. **I, J.** IBA1; **O, P.** CD3 and **U, V** Pax 5. **E, F.** Lungs at 14 DPI. **K, L.** IBA1; **Q, R.** CD3 and **W, X.** Pax 5. **A-F** HE. All images 200x. Abbreviations: RD = regular diet, HFHS = high-fat high-sugar, DPI = days post inoculation.

**Figure 6.**
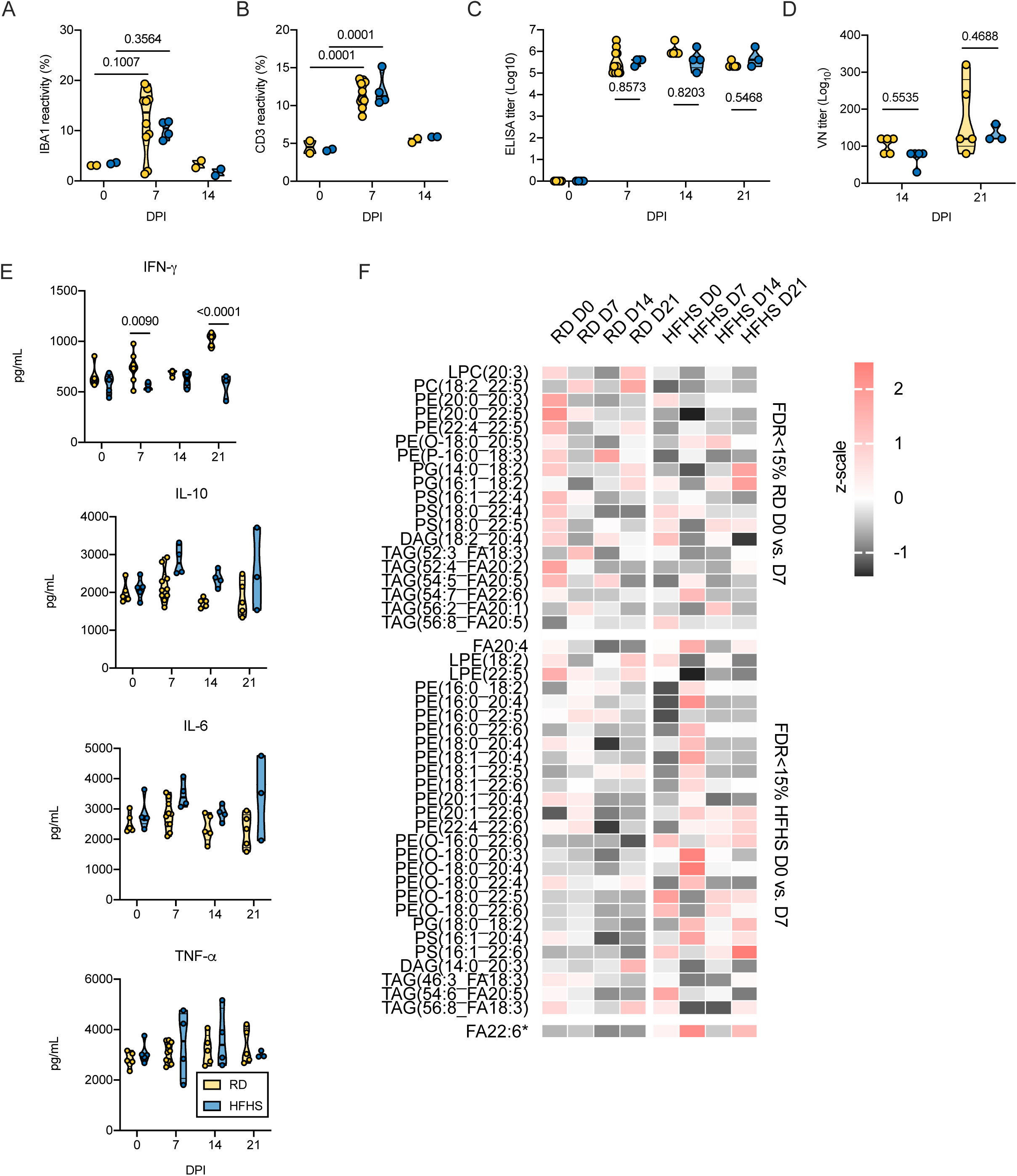
Disease manifestation is accompanied by prolonged viral shedding, systemic immune and metabolomic dysregulation after high-fat high-sugar diet. Animals were euthanized pre-challenge, at 7, 14, and 21 DPI with SARS-CoV-2 and serum and lung tissue collected for immune and lipid mediator analyses. Oropharyngeal swabs were taken to assess respiratory shedding **A.B.** Lung infiltration of T-cells (CD3) and macrophages (IBA1) was quantified by morphometric analysis. Truncated violin plots depicting median, quartiles and individuals, pre-challenge and 14 DPI: N = 2, 7 DPI: N = 10 (RD) / 4 (HFHS), ordinary two-way ANOVA, followed by Turkey’s multiple comparisons test. **C.** ELISA titers against spike protein of SARS-CoV-2 (lineage A) in serum obtained pre-challenge, at 7, 14, and 21 DPI. Truncated violin plots depicting median, quartiles and individuals, pre-challenge and 14 DPI: N = 5 (RD) / 4 (HFHS), 7 DPI: N = 10 (RD) / 4 (HFHS), 21 DPI: N = 5 (RD) / 3 (HFHS), ordinary two-way ANOVA, followed by Sidak’s multiple comparisons test. **D.** Virus neutralization titers against SARS-CoV-2 (lineage A) in serum obtained at 14 and 21 DPI. Truncated violin plots depicting median quartiles and individuals, 14 DPI: N = 5 (RD) / 4 (HFHS), 21 DPI: N = 5 (RD) / 3 (HFHS), ordinary two-way ANOVA, followed by Sidak’s multiple comparisons test. **E.F**. Viral load in oropharyngeal swabs measured in sgRNA copy number for RD and HFHS animals. Graphs show median, individual animals and 95% CI (shaded area). Dotted line = peak shedding. **G.** Area under the curve (AUC) analysis of virus shedding shown in E/F. Truncated violin plots depicting median quartiles and individuals, 21 DPI: N = 5 (RD) / 3 (HFHS), Mann-Whitney test. **H.** Serum levels (pg/mL) of INF-*γ*, TNF*α*-, IL-6 and IL-10 measured by ELISA from serum collected on 0, 7, 14, and 21 DPI. Truncated violin plots depicting median quartiles and individuals, pre-challenge/14 and 21 DPI: N = 5 (RD) / 4 (HFHS), 7 DPI: N = 10 (RD) / 4 (HFHS), ordinary two-way ANOVA, followed by Sidak’s multiple comparisons test. **I.** Lipid time-course heatmap: Changes in PUFA-containing serum lipids associated with an active SARS-CoV-2 infection as measured by LC-MS/MS. Autoscaled intensities are displayed for serum lipids species that were significantly changed between 0 and 7 DPI in either regular diet or HFHS diet hamsters with a false discovery rate of 15 % equating to p = 0.0256, 0.0193 for RD and HFHS, respectively. *FA22:6 (HFHS p = 0.0374) is displayed for comparison to clinical data despite not passing FDR filters. Abbreviations: TNF = tumor necrosis factor, IFN = interferon, IL = interleukin, RD = regular diet, HFHS = high-fat high-sugar, DPI = days post inoculation, sg = subgenomic, VN = virus neutralization. p-values are indicated were appropriate.

The humoral response to SARS-CoV-2 was not significantly impacted by diet regimen. Animals seroconverted at 7 DPI, as measured by anti-spike IgG ELISA (Fig 6 C, 7 DPI: N = 10 (RD) / 4 (HFHS), 14 DPI: N = 5 (RD) / 4 (HFHS), 21 DPI: N = 5 (RD) / 3 (HFHS), ordinary two-way ANOVA, followed by Tukey’s multiple comparisons test, p = 0.8573, p = 0.8203 and p = 0.5468, respectively). Neutralization of virus by sera collected at 14 and 21 DPI was compared to assess potential differences in affinity maturation and no significant difference was found (Fig 6 D, 14 DPI: N = 5 (RD) / 4 (HFHS), 21 DPI: N = 5 (RD) / 3 (HFHS), 14 DPI median = 120 / 80 and 21 DPI median = 120/120 reciprocal titer, ordinary two-way ANOVA, followed by Tukey’s multiple comparisons test, p = 0.5535 and p = 0.4688, respectively).

### Prolonged SARS-CoV-2 shedding, systemic immune and metabolomic dysregulation after high-fat high-sugar diet

The cytokine kinetics were analyzed in serum throughout the course of infection by ELISA. Serum samples were collected pre-challenge (0 DPI), on 7 DPI, 14 DPI and 21 DPI (Fig 6 E). Pro-inflammatory tumor necrosis factor (TNF)-*α*, interleukin (IL)-6, antiviral interferon (IFN)-*γ*, and (IL)-10 did not significantly differ between diet regimens pre-challenge. After infection, RD animals mounted a significant IFN-*γ* response which lasted into recovery (14 and 21 DPI), while no response was seen in HFHS animals (RD: N = 5/10, HFHS: N = 4, pre-challenge median = 629 / 618, 7 DPI median = 737.85 / 550.6, 14 DPI median = 702.3 / 623.55, 21 DPI median = 1042.3 / 609.8 pg/mL, ordinary two-way ANOVA, followed by Sidak’s multiple comparisons test, pre-challenge: p = 0.58157, 7 DPI: p = 0.0090, 14 DPI: p = 0.7373, 21 DPI p < 0.0001). In contrast, serum IL-6 trended higher in HFHS animals compared to RD animals at 7 DPI (median = 2795.5 (RD) / 2859.2 (HFHS) pg/mL). This trend toward higher IL-6 continued at 14 and 21 DPI. IL-10 levels trended higher in HFHS animals during the acute phase and remained elevated at 14 DPI (RD: N = 5/10, HFHS: N = 4, pre-challenge median = 1894.6 / 2131.5, 7 DPI median = 2071.75 / 2773.95, 14 DPI median = 1768.5 / 2354.35, 21 DPI median = 1733.7 / 2407.6 pg/mL ordinary two-way ANOVA, followed by Sidak’s multiple comparisons test, pre-challenge: p = 0.9933, 7 DPI: p = 0.0548, 14 DPI: p = 0.1408, 21 DPI p = 1259). TNF-*α* serum levels demonstrated an ambivalent pattern.

To examine compositional changes in the circulating lipidome over the course of infection, the lipidome was analyzed between 0 DPI and 7 DPI of infection. This analysis revealed distinct lipid dynamics in response to SARS-CoV-2 infection (Fig 6 F). RD animals displayed a serum lipid shift in response to infection consisting primarily of decreased levels of phospholipids with mixed representation of lipid classes and a distribution of long chain and polyunsaturated fatty acids (PUFA). HFHS serum displayed a more drastic pattern of lipid depletion and enrichment. Specifically, HFHS serum reflected a sharp enrichment of free polyunsaturated fatty acids (PUFA) and a combination of enrichment and depletion of PUFA containing phospholipids. This response peaked at 7 DPI and began to return to homeostasis by 14 DPI, though certain lipid patterns were carried out until 21 DPI.

## Discussion

The development of animal models that faithfully recapitulate certain aspects of human disease remains a top priority in SARS-CoV-2 research. Healthy Syrian hamsters develop mild to moderate disease similar to the majority of human cases; however, they do not exhibit the more severe respiratory disease seen in humans with comorbidities such as obesity, diabetes, or other chronic illness (Araújo et al., 2019; Hussain et al., 2020; Korakas et al., 2020). Thus, we developed an experimental infection model of hamsters exclusively fed a high-fat high-sugar diet to model the impact of Western Diet on COVID-19 severity. In the Syrian hamster, this diet caused diet-induced morbidity, led to increased weight gain during adolescence, and ultimately led to in increased glucose tolerance, systemic hyperlipidemia, increased total cholesterol and a liver pathology reminiscent of a NAFLD-like phenotype. The lack of net weight gain in this model may present a means of decoupling liver associated pathologies such as NAFLD from obesity-associated disease more broadly. In humans NAFLD is predominantly a consequence of obesity and frequently associated also with other comorbidities as well (Sanyal, 2019). In the context of COVID-19, NAFLD is associated with increased hospitalization and disease severity (Bramante et al., 2020).

The morbidity observed in the absence of infection in the HFHS group should be considered in future studies utilizing this model. In particular, this feature of the model may make survival-based studies difficult. Human clinical studies of COVID-19 are plagued by this same difficulty in quantifying the contribution of infection and the associated comorbidities to the eventual cause of death. If appropriately controlled for in this model the relative contribution to death from the infection and the comorbidities can be quantified. We observed that male hamsters on a HFHS diet demonstrated delayed lower and upper respiratory tract clearance after infection with SARS-CoV-2, which was accompanied by more severe disease presentation. Our data is in agreement with findings in mice, which have reported enhanced morbidity in aged and diabetic obese mice in a mouse-adapted SARS-CoV-2 model (Rathnasinghe et al., 2021). Conversely, we also observed increased weight loss, pathology, delayed lung recovery and influx of immune cells into the lung in a subset of hamsters fed a regular diet as compared to what has been shown in younger animals (Chan et al., 2020; Rosenke et al., 2020). This is likely due to the increased age of the animals used in this study (Osterrieder et al., 2020). Previously, lung function analysis after SARS-CoV-2 infection in a rodent model has only been demonstrated in ACE2 mice (Winkler et al., 2020). While not significantly different between the diet groups, we performed functional lung analysis for the first time in the Syrian hamster after SARS-CoV-2 infection and demonstrated that this model also recapitulates increased total airway resistance and decreased inspiratory capacity. This suggests that the Syrian hamster, besides recapitulating lung pathology, may also be a useful model for mechanistic studies of the respiratory parameters affected by COVID-19. Importantly, the HFHS Syrian hamster model presented here recapitulated two key mediators of severe human COVID-19. One unique feature of the cytokine profile in human disease is the elevation of IL-6 and IL-10, which have been indicated as causes of increased pathology (Chen et al., 2020; Dhar et al., 2020; Lu et al., 2021; Wang et al., 2020). In line with this, in HFHS animals we observed trending increases in serum IL-10 and IL-6 levels after infection. Secondly, in response to infection, HFHS animals showed a more severe response in their serum lipids at 7 DPI compared to RD animals. The lipids that dominated this response were free-PUFAs and PUFA-containing phosphatidylethanolamine (PE). In addition, we saw mixed increase and decrease of PUFA-containing plasmalogens and triacylglycerols. The metabolic comorbidities associated with severe COVID-19 were previously shown to correlate with specific mobilization of serum lipids in a human cohort (Schwarz et al., 2020). Specifically, disease severity, defined by ICU admittance, was shown to be associated with increased free PUFAs and PUFA-containing phosphatidylethanolamine, as well as a decrease of PUFA-containing phosphatidylcholine and plasmalogen, compared to non-ICU hospitalized patients. These imbalances were reflected in the circulating milieu of immune-active, PUFA-derived lipid mediators in these patients. The lipid pattern findings in the Syrian hamster model suggest that these serum lipid changes are dependent on preexisting serum hyperlipidemia and stimulated by infection with SARS-CoV-2. Despite the lack of obesity in these animals, the matching of clinical SARS-CoV-2-associated lipid patterns and cytokine profile in this model supports its utility in examining lipid and inflammation dynamics associated immune dysregulation during infection.

Of note, this did not seem to adversely affect the humoral immune response while viral titers in oropharyngeal swabs and lung tissues suggested delayed clearance in the HFHS group. This may indicate that other immune pathways were disproportionately affected, but further investigations would be necessary to draw concrete conclusions.

Taking the limitations of the model into account, our data further suggests the possible suitability of the Syrian hamster model to assess immunomodulatory therapies. While dietary advice for those suffering from metabolic diseases is proposed to reduce burden of severe COVID-19 (Demasi, 2021), it remains doubtful if any change in diet can impact disease outcome favorably after infection has occurred. Targeted immunomodulatory therapies, such as anti-IL-6 therapies, may be more efficient (Zhong et al., 2020). The Syrian hamster model may also be applied to further studies of selected aspects of NAFLD, which the model recapitulates. This model seems to present with an absences or limited amount of liver fibrosis; further work is needed to demonstrate how faithfully it assesses the direct effect of liver fibrosis on acute disease. However, it may be useful to assess long term post-COVID-19 NAFLD, to document further deterioration of liver damage (Portincasa et al., 2020) and the relation to infection sequelae.

## Acknowledgements

The authors would like to thank Kwe Claude Yinda and Robert Fisher for technical support, the Rocky Mountain Veterinary branch, including Marissa Woods, Nicki Arndt, Amanda Weidow, Linda Couey and Brian Mosbrucker for assistance with high containment husbandry, Tina Thomas, and Rebecca Rosenke for assistance with histology and Danielle Hopkins for flexiVent support and Kathryn Willebrand and Taylor Saturday for assistance with manuscript editing. This research was supported by the Intramural Research Program of the National Institute of Allergy and Infectious Diseases (NIAID), National Institutes of Health (NIH).

## Author Contributions

Conceptualization, J.R.P.;

Methodology, J.R.P., D.R.A., and B.S.;

Investigation, J.R.P., D.R.A.,B.S, J.E.S., V.A.A., M.G.H., J.N.P., D.S, K.A.S., I.L., B.J.S., C.S.;

Writing – Original Draft, J.R.P., D.R.A., B.S., C.S.;

Writing – Review & Editing, J.R.P., D.R.A. and V.J.M.;

Funding Acquisition, VJ..M.;

Resources, V.A.A., B.S., B.J.S. and C.M.B.;

Supervision, V.J.M.

## Methods

### Ethics statement

Approval of animal experiments was obtained from the Institutional Animal Care and Use Committee of the Rocky Mountain Laboratories. Performance of experiments was done following the guidelines and basic principles in the United States Public Health Service Policy on Humane Care and Use of Laboratory Animals and the Guide for the Care and Use of Laboratory Animals. Work with infectious SARS-CoV-2 strains under BSL3 conditions was approved by the Institutional Biosafety Committee (IBC). Inactivation and removal of samples from high containment was performed per IBC-approved standard operating procedures (Haddock et al., 2021).

### Virus and cells

SARS-CoV-2 strain nCoV-WA1-2020 (MN985325.1) was provided by CDC, Atlanta, USA. Virus propagation was performed in VeroE6 cells in DMEM supplemented with 2% foetal bovine serum (FBS), 2 mM L-glutamine, 100 U/mL penicillin and 100 μg/mL streptomycin. VeroE6 cells were maintained in DMEM supplemented with 10% FBS, 2 mM L-glutamine, 100 U/mL penicillin and 100 μg/mL streptomycin D10. Virus stock was 100% identical to the initial sequence (MN985325.1) and no contaminants were detected.

### High-fat high-sugar diet

Four to six-week-old male Syrian Golden hamsters (ENVIGO) were randomly assigned to either regular rodent chow (Teklad Global 16% Protein Rodent Diet, Envigo) or a HFHS diet for 16 weeks (Purina Chow #5001 with 11.5% Corn Oil, 11.5% Coconut Oil, 0.5% Cholesterol, 0.25% Deoxycholic Acid, and 10% Fructose: Dyets Inc., Dyet#615088). Pre-challenge oral glucose tests were performed on all animals. Five animals from each diet group were euthanized after the 16 wks for collection of pre-challenge tissue samples and weights. For each diet group, 5 animals were randomly designated for flexiVent calibration and excluded from further analysis. Three animals in the HFHS regimen were euthanized throughout the 16-week diet regimen due to secondary morbidities and were not included in analyses. Pre-challenge, an additional 5 animals in the RD group and additional 8 animals in the HFHS group were excluded from the study due to experimental reasons, and one animal in the HFHS group due to secondary morbidities.

### Assessment of glucose tolerance

An oral glucose tolerance test (OGTT) was performed after 16 weeks of diet manipulation (Dalbøge et al., 2015). Hamsters were fasted for 16 h overnight preceding the OGTT. An oral glucose load (2 g/kg glucose) was administered. Blood samples were collected from the retroorbital sinus using capillary tube at 0-, 30-, 60-, and 120-min post glucose administration. Blood glucose was measured using the AlphaTRAK blood glucose monitoring system (Zoetis), calibrated for cats. Serum was separated and used for measurement of insulin. Insulin was measured using the rat/mouse insulin ELISA kit from Millipore (EZRMI-13K), according to the manufacturer’s instructions (Wang et al., 2001).

### Lipidomics

Blood lipids were assessed for a subset of animals (N= 8-10) after 16 weeks of diet. 200 µL blood was collected and were measured using the Piccolo® Lipid Panel Plus for humans (Abraxis) according to the manufacturer’s instruction.

### Next-generation sequencing of liver mRNA

Frozen tissues were pulverized in 1 mL of Trizol (Thermofisher Scientific), 200 µL of 1-Bromo-3-chloropropane (MilliporeSigma) was added, samples mixed, and centrifuged at 16,000 x *g* for 15 min at 4 °C. RNA containing aqueous phase of 600 µL was collected from each sample and passed through Qiashredder column (Qiagen) at 21,000 x g for 2 min to homogenize any remaining genomic DNA in the aqueous phase. Aqueous phase was combined with 600 µL of RLT lysis buffer (Qiagen, Valencia, CA) with 1% beta mercaptoethanol (MilliporeSigma) and RNA was extracted using Qiagen AllPrep DNA/RNA 96-well system. An additional on-column DNase-1 treatment was performed during RNA extraction. RNA was quantitated by spectrophotometry and yield ranged from 0.4 to 17.8 µg. One hundred nanograms of RNA was used as input for rRNA depletion and NGS library preparation following the Illumina Stranded Total RNA Prep Ligation with Ribo-Zero Plus workflow (Illumina). The NGS libraries were prepared, amplified for 13 cycles, AMPureXP bead (Beckman Coulter) purified using 0.95X beads, assessed on a BioAnalyzer DNA1000 chip (Agilent Technologies) and quantified using the Kapa Quantification Kit for Illumina Sequencing (Roche). Amplified libraries were pooled at equal molar amounts and sequenced on a NextSeq (Illumina) using two High Output 150 cycle chemistry kits. Raw fastq reads were trimmed of Illumina adapter sequences using cutadapt version 1.12 and then trimmed and filtered for quality using the FASTX-Toolkit (Hannon Lab). Remaining reads were aligned to the *Mesocricetus auratus* genome assembly version 1.0 using Hisat2 (Kim et al., 2015). Reads mapping to genes were counted using htseq-count (Anders et al., 2015). Differential expression analysis was performed using the Bioconductor package DESeq2 (Love et al., 2014). Pathway analysis was performed using Ingenuity Pathway Analysis (QIAGEN) and gene clustering was performed using Partek Genomics Suite (Partek Inc.). Samples with too low quality were removed from the analysis (Sup Table 1).

### Next-generation sequencing of virus

For sequencing from viral stocks, sequencing libraries were prepared using Stranded Total RNA Prep Ligation with Ribo-Zero Plus kit per manufacturer’s protocol (Illumina) and sequenced on an Illumina MiSeq at 2 x 150 base pair reads. For sequencing from swab and lung tissue, total RNA was depleted of ribosomal RNA using the Ribo-Zero Gold rRNA Removal kit (Illumina). Sequencing libraries were constructed using the KAPA RNA HyperPrep kit following manufacturer’s protocol (Roche Sequencing Solutions). To enrich for SARS-CoV-2 sequence, libraries were hybridized to myBaits Expert Virus biotinylated oligonucleotide baits following the manufacturer’s manual, version 4.01 (Arbor Biosciences). Enriched libraries were sequenced on the Illumina MiSeq instrument as paired-end 2 × 150 base pair reads. Raw fastq reads were trimmed of Illumina adapter sequences using cutadapt version 1.1227 and then trimmed and filtered for quality using the FASTX-Toolkit (Hannon Lab, CSHL). Remaining reads were mapped to the SARS-CoV-2 2019-nCoV/USA-WA1/2020 genome (MN985325.1) using Bowtie2 version 2.2.928 with parameters --local --no-mixed -X 1500. PCR duplicates were removed using picard MarkDuplicates (Broad Institute) and variants were called using GATK HaplotypeCaller version 4.1.2.029 with parameter -ploidy 2. Variants were filtered for QUAL > 500 and DP > 20 using bcftools.

### Inoculation experiments

After 16 weeks, animals were then inoculated intranasally (I.N.) under isoflurane anaesthesia. I.N. inoculation was performed with 40 µL sterile Dulbecco’s Modified Eagle Medium (DMEM) containing 8×10^4^ TCID_50_ SARS-CoV-2. A subset of animals (N= 4-10) were euthanized, and serum and tissues were collected at pre-challenge (0 DPI), 4, 7, 14, and 21 DPI. Hamsters were weighted daily, and oropharyngeal swabs (21 DPI animals only) were taken daily until day 7 and then thrice a week. Swabs were collected in 1 mL DMEM with 200 U/mL penicillin and 200 µg/mL streptomycin. Hamsters were observed daily for clinical signs of disease.

### Lung function analyses

Lung function assessment was performed on pre-challenge, 7, 14, and 21 DPI. Hamsters were anesthetized with a combination of inhalant isoflurane and ketamine/xylazine intraperitoneally. After animals reached a surgical plane of anaesthesia a terminal tracheostomy was performed as previously described (McGovern TK JOVE 2013). Briefly, a cannula was introduced into the trachea, secured with suture, and the animal underwent the forced oscillation technique (FOT) using a flexiVent (SCIREZ, Inc.). Animals were kept at a consistent surgical plane of anesthesia to the point of not resisting the FOT procedure. Animals were immediately euthanized while deeply anesthetized after FOT was completed; the surgical procedure was terminal.

### Histopathology and immunohistochemistry

Necropsies and tissue sampling were performed according to IBC-approved protocols. Tissues were fixed for a minimum of 7 days in 10% neutral buffered formalin with 2 changes. Tissues were placed in cassettes and processed with a Sakura VIP-6 Tissue Tek, on a 12-hour automated schedule, using a graded series of ethanol, xylene, and ParaPlast Extra. Prior to staining, embedded tissues were sectioned at 5 µm and dried overnight at 42°C. Using GenScript U864YFA140-4/CB2093 NP-1 (1:1000) specific anti-CoV immunoreactivity, CD3 (Predilute) (Roche Tissue Diagnostics #790-4341), and PAX5 (1:500) (Novus Biologicals #NBP2-38790) were detected using the Vector Laboratories ImPress VR anti-rabbit IgG polymer (# MP-6401) as the secondary antibody. Iba-1 (1:500) (abcam #ab5076) was detected using Roche Tissue Diagnostics OmniMap anti-goat multimer (#760-4647) as the secondary antibody. The tissues were stained using the Discovery Ultra automated stainer (Ventana Medical Systems) with a ChromoMap DAB kit Roche Tissue Diagnostics (#760-159).

### Morphometric analysis

IHC stained tissue slides were scanned with an Aperio ScanScope XT (Aperio Technologies, Inc.) and analyzed using the ImageScope Positive Pixel Count algorithm (version 9.1). The default parameters of the Positive Pixel Count (hue of 0.1 and width of 0.5) detected antigen adequately.

### Viral RNA detection

Swabs from hamsters were collected as described above. Cage and bedding material was sampled with prewetted swabs in 1 mL of DMEM supplemented with 200 U/mL penicillin and 200 μg/mL streptomycin. Then, 140 µL was utilized for RNA extraction using the QIAamp Viral RNA Kit (Qiagen) using QIAcube HT automated system (Qiagen) according to the manufacturer’s instructions with an elution volume of 150 µL. Sub-genomic (sg) viral RNA and genomic (g) was detected by qRT-PCR (Corman et al., 2020a; Corman et al., 2020b). Five μL RNA was tested with TaqMan™ Fast Virus One-Step Master Mix (Applied Biosystems) using QuantStudio 6 Flex Real-Time PCR System (Applied Biosystems) according to instructions of the manufacturer. Ten-fold dilutions of SARS-CoV-2 standards with known copy numbers were used to construct a standard curve and calculate copy numbers/mL.

### Viral titration

Viable virus in tissue samples was determined as previously described (van Doremalen et al., 2017). In brief, lung tissue samples were weighted, then homogenized in 1 mL of DMEM2. VeroE6 cells were inoculated with ten-fold serial dilutions of tissue homogenate, spun at 1000 rpm for 1 h at 37 °C, the first dilutions washed with PBS and with DMEM2. Cells were incubated with tissue homogenate for 6 days at 37 °C, 5% CO_2_, then scored for cytopathic effect. TCID_50_ was calculated by the method of Spearman-Karber and adjusted for tissue weight.

### Serology

Serum samples were inactivated with γ-irradiation (2 mRad) and analyzed as previously described (Yinda et al., 2020). In brief, maxisorp plates (Nunc) were coated with 50 ng spike protein (generated in-house) per well and incubated overnight at 4 °C. After blocking with casein in phosphate buffered saline (PBS) (ThermoFisher) for 1 h at room temperature (RT), serially diluted 2-fold serum samples (duplicate, in blocking buffer) were incubated for 1 h at RT. Spike-specific antibodies were detected with goat anti-hamster IgG Fc (horseradish peroxidase (HRP)-conjugated, Abcam) for 1 h at RT and visualized with KPL TMB 2-component peroxidase substrate kit (SeraCare, 5120-0047). The reaction was stopped with KPL stop solution (Seracare) and read at 450 nm. Plates were washed 3 to 5 x with PBS-T (0.1 % Tween) for each wash. The threshold for positivity was calculated as the average plus 3 x the standard deviation of negative control hamster sera.

### Cytokine analysis

Cytokine concentrations were determined using a commercial hamster ELISA kit for TNF-*α*, INF-*γ*, IL-6, IL-4, and IL-10 available at antibodies.com, according to the manufacturer’s instructions (antibodies.com; A74292, A74590, A74291, A74027, A75096). Samples were pre-diluted 1:10.

### Serum lipid analysis

For abundance analysis of serum lipids signals were filtered using a 50 % miss value cut off and applying a raw intensity cutoff appropriate to the noise level of each class of lipids. Signals were then normalized to internal deuterated SPLASH® LIPIDOMIX® Mass Spec Standard (Avanti Polar Lipids). For compositional analysis of the serum, bulk lipid datasets were further filtered using a 30 % QC coefficient of variance cut off prior to normalizing by the total signal sum. All univariate and multivariate analysis was performed using GraphPad Prism or MarkerView (AB Sciex). All parallel univariate analysis was subjected to a Benjamini-Hochberg correction using a false discovery rate of 15 %.

### Statistical analysis

All graphs were designed in GraphPad Prism software (version 8.0.1; GraphPad Software). Significance test were performed as indicated where appropriate. Statistical significance levels were determined as follows: ns = p > 0.05; * = p ≤ 0.05; ** = p ≤ 0.01; *** = p ≤ 0.001; **** = p ≤ 0.0001.

## Supplemental Material

**Supplemental Table 1:** Liver marker profile in serum of regular diet (RD) and high-fat high-sugar diet (HFHS) after 16 weeks. Quantitative determination of total cholesterol (CHOL), high-density lipoprotein cholesterol (HDL), triglycerides (TRIG), alanine aminotransferase (ALT), aspartate aminotransferase (AST), and glucose (GLU) in heparinized whole blood. From the CHOL, HDL and TRIG determinations, low-density lipoprotein cholesterol (LDL), very low-density lipoprotein cholesterol (VLDL), non-HDL cholesterol, and a total cholesterol/high-density lipoprotein cholesterol ratio (TC/H) was calculated. ∼∼ + could not be calculated, LIP = not detectable due to lipid interference.

| Animal ID | Chol<br>mg/dl | HDL<br>mg/dl | Trig<br>mg/dl | ALT<br>U/L | AST<br>U/L | GLU<br>mg/dl | nHDL<br>Lc<br>mg/dl | TC/<br>H | LDL<br>mg/dl | VLDL<br>mg/dl | LIP |
| --- | --- | --- | --- | --- | --- | --- | --- | --- | --- | --- | --- |
| HFHS.1 | 360 | LIP | ~~~ | 124 | 78 | 120 | ~~~ | ~~~ | LIP | LIP | 3 |
| HFHS.2 | >520 | HEM | HEM | 224 | HEM | LIP | ~~~ | ~~~ | ~~~ | ~~~ | 3 |
| HFHS.3 | ~~~ | HEM | ~~~ | 140 | HEM | LIP | ~~~ | ~~~ | ~~~ | ~~~ | 3 |
| HFHS.4 | >520 | HEM | HEM | 190 | HEM | LIP | ~~~ | ~~~ | ~~~ | ~~~ | 3 |
| HFHS.5 | 380 | LIP | >500 | 117 | 82 | 114 | ~~~ | ~~~ | LIP | LIP | 3 |
| HFHS.6 | 263 | LIP | >500 | 152 | 86 | 110 | ~~~ | ~~~ | LIP | LIP | 3 |
| HFHS.7 | 384 | ~~~ | ~~~ | 188 | 147 | LIP | ~~~ | ~~~ | ~~~ | ~~~ | 3 |
| HFHS.8 | 331 | LIP | >500 | 114 | 84 | 59 | ~~~ | ~~~ | LIP | LIP | 2 |
| Regular.1 | 83 | 49 | 212 | 62 | 76 | 113 | 34c | 1.7c | 0 | 42c | 1 |
| Regular.2 | 52 | 31 | 168 | 100 | 152 | 151 | 21c | 1.7c | 0 | 34c | 0 |
| Regular.3 | 98 | 68 | 248 | 67 | 79 | 72 | 30c | 1.4c | 0 | 50c | 0 |
| Regular.4 | 111 | 92 | 252 | 87 | 87 | 79 | 19c | 1.2c | ~~~ | 50c | 1 |
| Regular.5 | 74 | 43 | 184 | 73 | 96 | 98 | 31c | 1.7c | 0 | 37c | 1 |
| Regular.6 | 59 | 32 | 201 | 87 | 113 | 119 | 27c | 1.8c | 0 | 40c | 1 |
| Regular.7 | 43 | 23 | 204 | 148 | 127 | 101 | 20c | 1.8c | ~~~ | 41c | 0 |
| Regular.8 | 71 | 42 | 198 | 80 | 88 | 85 | 29c | 1.7c | 0 | 40c | 0 |
| Regular.9 | 64 | 46 | 232 | 142 | 157 | 88 | 18c | 1.4c | ~~~ | 46c | 0 |
| Regular.10 | 49 | 28 | 251 | 123 | 166 | 99 | 21c | 1.7c | ~~~ | 50c | 0 |

**Supplemental Table 2:** Up-and down-regulated pathways in livers pre-challenge organized by disease and function.

| **Diseases or Functions**<br>**Annotation** | **p-value** | **Predicted**<br>**Activation State** | **Activation**<br>**z-score** | **#**<br>**Molecules** |
| --- | --- | --- | --- | --- |
| Development of genitourinary system | 2.07E-11 | Increased | 2.114 | 160 |
| Internalization of cells | 1.99E-11 | Increased | 5.333 | 63 |
| Abnormal bone density | 1.99E-11 | Decreased | -2.297 | 50 |
| Phagocytosis of blood cells | 1.89E-11 | Increased | 4.815 | 52 |
| Adhesion of lymphocytes | 1.54E-11 | Increased | 3.747 | 34 |
| Interaction of T lymphocytes | 1.36E-11 | Increased | 3.906 | 39 |
| Adhesion of lymphatic system cells | 1.33E-11 | Increased | 3.839 | 35 |
| Cell movement of macrophages | 1.32E-11 | Increased | 4.236 | 65 |
| Adhesion of tumor cell lines | 9.66E-12 | Increased | 2.42 | 68 |
| Pancreatobiliary tumor | 9.07E-12 | Increased | 2.146 | 402 |
| Size of body | 9.03E-12 | Increased | 2.966 | 126 |
| Cell cycle progression | 8.08E-12 | Increased | 2.341 | 171 |
| Binding of T lymphocytes | 7.98E-12 | Increased | 3.521 | 37 |
| Interaction of lymphocytes | 5.70E-12 | Increased | 3.973 | 45 |
| Pancreatic lesion | 5.57E-12 | Increased | 2.114 | 358 |
| Migration of neutrophils | 5.56E-12 | Increased | 3.941 | 38 |
| Response of myeloid leukocytes | 5.43E-12 | Increased | 2.565 | 34 |
| Chemotaxis of neutrophils | 5.10E-12 | Increased | 2.328 | 42 |
| Migration of granulocytes | 4.82E-12 | Increased | 3.2 | 43 |
| Cell-cell contact | 4.15E-12 | Increased | 2.864 | 136 |
| Development of head | 3.98E-12 | Increased | 3.325 | 164 |
| Quantity of metal ion | 3.75E-12 | Increased | 2.912 | 85 |
| Aggregation of blood platelets | 3.23E-12 | Increased | 3.084 | 49 |
| Immune response of antigen presenting cells | 3.23E-12 | Increased | 3.968 | 49 |
| Binding of lymphatic system cells | 2.58E-12 | Increased | 3.801 | 45 |
| Transmigration of leukocytes | 2.57E-12 | Increased | 2.603 | 42 |
| Binding of lymphocytes | 2.47E-12 | Increased | 3.619 | 43 |
| Transport of molecule | 2.41E-12 | Increased | 2.741 | 246 |
| Malignant connective or soft tissue neoplasm | 2.36E-12 | Increased | 2.079 | 203 |
| Response of antigen presenting cells | 1.93E-12 | Increased | 4.049 | 52 |
| Recruitment of macrophages | 1.92E-12 | Increased | 2.61 | 33 |
| Engulfment of cells | 1.87E-12 | Increased | 4.959 | 99 |
| Phagocytosis | 1.52E-12 | Increased | 5.163 | 81 |
| Homing of neutrophils | 1.47E-12 | Increased | 2.328 | 43 |
| Phagocytosis of cells | 1.45E-12 | Increased | 5.586 | 75 |
| Binding of endothelial cells | 1.13E-12 | Increased | 2.496 | 49 |
| Transmigration of cells | 1.13E-12 | Increased | 2.844 | 49 |
| Cell movement of cancer cells | 8.81E-13 | Increased | 2.68 | 41 |
| Interaction of endothelial cells | 8.80E-13 | Increased | 2.389 | 50 |
| Cellular infiltration by myeloid cells | 8.74E-13 | Increased | 2.023 | 72 |
| Inflammation of respiratory system component | 7.19E-13 | Increased | 2.017 | 105 |
| Binding of lymphoid cells | 7.09E-13 | Increased | 3.713 | 44 |
| Activation of antigen presenting cells | 6.15E-13 | Increased | 3.438 | 70 |
| Homeostasis of blood cells | 5.80E-13 | Increased | 3.757 | 111 |
| Activation of myeloid cells | 4.67E-13 | Increased | 3.734 | 74 |
| Degranulation of phagocytes | 4.12E-13 | Increased | 3.533 | 90 |
| Activation of phagocytes | 3.71E-13 | Increased | 3.826 | 79 |
| Engulfment of myeloid cells | 3.71E-13 | Increased | 4.655 | 47 |
| Degranulation of myeloid cells | 3.58E-13 | Increased | 3.647 | 91 |
| Synthesis of reactive oxygen species | 2.88E-13 | Increased | 4.241 | 99 |
| Homeostasis of leukocytes | 2.75E-13 | Increased | 3.757 | 110 |
| Malignant neoplasm of retroperitoneum | 2.37E-13 | Increased | 2.021 | 419 |
| Quantity of Ca<sup>2+</sup> | 2.14E-13 | Increased | 2.613 | 82 |
| Engulfment of leukocytes | 2.00E-13 | Increased | 4.233 | 51 |
| Recruitment of myeloid cells | 1.80E-13 | Increased | 4.402 | 64 |
| T cell development | 1.58E-13 | Increased | 3.822 | 104 |
| Degranulation of leukocytes | 1.56E-13 | Increased | 3.4 | 95 |
| Upper gastrointestinal tract tumor | 1.49E-13 | Increased | 2.236 | 581 |
| Invasion of cells | 1.48E-13 | Increased | 4.617 | 184 |
| Engulfment of phagocytes | 1.43E-13 | Increased | 4.266 | 49 |
| Cell movement of tumor cell lines | 1.37E-13 | Increased | 4.127 | 185 |
| Metabolism of reactive oxygen species | 1.34E-13 | Increased | 4.381 | 104 |
| Growth of connective tissue | 1.31E-13 | Increased | 2.459 | 123 |
| Amyloidosis | 1.30E-13 | Decreased | -2.433 | 116 |
| Activation of mononuclear leukocytes | 1.19E-13 | Increased | 3.257 | 95 |
| Upper gastrointestinal tract cancer | 1.13E-13 | Increased | 2 | 580 |
| Lymphopoiesis | 9.10E-14 | Increased | 3.693 | 123 |
| Adhesion of mononuclear leukocytes | 7.27E-14 | Increased | 3.616 | 42 |
| Quantity of immunoglobulin | 6.65E-14 | Increased | 3.038 | 63 |
| Production of antibody | 6.01E-14 | Increased | 3.204 | 66 |
| Cell viability | 5.81E-14 | Increased | 4.969 | 235 |
| Phagocytosis of phagocytes | 4.62E-14 | Increased | 4.14 | 46 |
| Phagocytosis of leukocytes | 4.60E-14 | Increased | 4.229 | 47 |
| Recruitment of antigen presenting cells | 4.55E-14 | Increased | 3.038 | 38 |
| Phagocytosis of myeloid cells | 3.83E-14 | Increased | 4.396 | 46 |
| Activation of lymphocytes | 3.64E-14 | Increased | 3.171 | 93 |
| Cell movement of T lymphocytes | 3.42E-14 | Increased | 3.59 | 63 |
| Recruitment of phagocytes | 3.01E-14 | Increased | 4.827 | 62 |
| Cell survival | 1.69E-14 | Increased | 4.795 | 247 |
| Activation of lymphoid cells | 1.50E-14 | Increased | 3.25 | 94 |
| Migration of myeloid cells | 1.43E-14 | Increased | 3.972 | 54 |
| Production of protein | 1.43E-14 | Increased | 3.73 | 70 |
| Quantity of B lymphocytes | 1.29E-14 | Increased | 2.549 | 81 |
| Activation of lymphatic system cells | 1.09E-14 | Increased | 3.088 | 95 |
| Metastasis | 1.03E-14 | Increased | 3.523 | 181 |
| Interaction of mononuclear leukocytes | 9.60E-15 | Increased | 3.685 | 55 |
| Growth of epithelial tissue | 7.66E-15 | Increased | 2.312 | 135 |
| Quantity of T lymphocytes | 7.33E-15 | Increased | 4.155 | 111 |
| T cell homeostasis | 7.05E-15 | Increased | 3.667 | 109 |
| Hematopoiesis of mononuclear leukocytes | 6.16E-15 | Increased | 3.759 | 132 |
| Migration of antigen presenting cells | 4.15E-15 | Increased | 3.735 | 51 |
| Immediate hypersensitivity | 4.06E-15 | Increased | 2.362 | 77 |
| Response of myeloid cells | 3.82E-15 | Increased | 4.283 | 60 |
| Immune response of myeloid cells | 2.85E-15 | Increased | 4.069 | 56 |
| Non-colon gastrointestinal cancer | 2.59E-15 | Increased | 2 | 612 |
| Aggregation of cells | 2.51E-15 | Increased | 3.773 | 77 |
| Extraadrenal retroperitoneal tumor | 2.51E-15 | Increased | 2.58 | 463 |
| Chemotaxis of myeloid cells | 2.36E-15 | Increased | 3.493 | 70 |
| Leukopoiesis | 2.22E-15 | Increased | 4.158 | 149 |
| Aggregation of blood cells | 1.69E-15 | Increased | 3.56 | 62 |
| Differentiation of mononuclear leukocytes | 1.67E-15 | Increased | 3.796 | 134 |
| Hereditary connective tissue disorder | 1.43E-15 | Decreased | -3.259 | 128 |
| Binding of mononuclear leukocytes | 1.33E-15 | Increased | 3.331 | 54 |
| Advanced malignant tumor | 8.44E-16 | Increased | 3.434 | 197 |
| Connective tissue tumor | 7.45E-16 | Increased | 2.495 | 222 |
| Advanced stage tumor | 5.25E-16 | Increased | 3.434 | 198 |
| Chemotaxis of phagocytes | 4.47E-16 | Increased | 3.987 | 73 |
| Recruitment of blood cells | 4.05E-16 | Increased | 4.395 | 79 |
| Connective or soft tissue tumor | 3.53E-16 | Increased | 2.525 | 249 |
| Degranulation of cells | 3.26E-16 | Increased | 3.573 | 115 |
| Recruitment of leukocytes | 2.92E-16 | Increased | 4.208 | 78 |
| Chemotaxis of leukocytes | 2.63E-16 | Increased | 4.146 | 84 |
| Immune response of phagocytes | 2.22E-16 | Increased | 4.017 | 62 |
| Growth of tumor | 2.05E-16 | Increased | 4.005 | 179 |
| Degranulation | 1.09E-16 | Increased | 3.507 | 117 |
| Inflammation of joint | 1.07E-16 | Increased | 3.397 | 178 |
| Response of phagocytes | 9.69E-17 | Increased | 4.268 | 66 |
| Chemotaxis of blood cells | 9.45E-17 | Increased | 4.147 | 85 |
| Recruitment of cells | 8.37E-17 | Increased | 4.692 | 85 |
| T cell migration | 7.55E-17 | Increased | 4.271 | 73 |
| Immune response of leukocytes | 7.50E-17 | Increased | 4.727 | 80 |
| Hypersensitive reaction | 6.65E-17 | Increased | 3.831 | 97 |
| Vasculogenesis | 5.92E-17 | Increased | 3.35 | 158 |
| Angiogenesis | 5.56E-17 | Increased | 4.04 | 184 |
| Development of vasculature | 4.93E-17 | Increased | 3.955 | 198 |
| Binding of tumor cell lines | 4.04E-17 | Increased | 2.518 | 93 |
| Homing of leukocytes | 3.51E-17 | Increased | 4.4 | 89 |
| Cell movement of antigen presenting cells | 2.39E-17 | Increased | 4.024 | 91 |
| Binding of myeloid cells | 2.30E-17 | Increased | 3.053 | 60 |
| Cellular homeostasis | 2.19E-17 | Increased | 4.676 | 269 |
| Experimental autoimmune encephalomyelitis | 1.82E-17 | Increased | 3.58 | 86 |
| Homing of blood cells | 1.80E-17 | Increased | 4.405 | 90 |
| Interaction of tumor cell lines | 1.57E-17 | Increased | 2.283 | 96 |
| Microtubule dynamics | 6.51E-18 | Increased | 3.927 | 216 |
| Cell movement of granulocytes | 5.63E-18 | Increased | 3.593 | 93 |
| Cell movement of lymphatic system cells | 2.90E-18 | Increased | 4.135 | 101 |
| Organization of cytoplasm | 2.90E-18 | Increased | 4.437 | 262 |
| Cell movement of neutrophils | 2.55E-18 | Increased | 3.635 | 83 |
| Encephalitis | 2.39E-18 | Increased | 2.541 | 95 |
| Immune response of cells | 1.57E-18 | Increased | 4.854 | 130 |
| Cell movement of lymphocytes | 1.14E-18 | Increased | 4.249 | 100 |
| Allergy | 5.20E-19 | Increased | 3.26 | 98 |
| Rheumatic Disease | 4.02E-19 | Increased | 3.322 | 223 |
| Organismal death | 3.96E-19 | Decreased | -4.3 | 362 |
| Morbidity or mortality | 2.66E-19 | Decreased | -4.307 | 366 |
| Cell proliferation of T lymphocytes | 2.10E-19 | Increased | 2.415 | 127 |
| Development of body trunk | 1.36E-19 | Increased | 3.651 | 215 |
| Binding of professional phagocytic cells | 1.35E-19 | Increased | 2.925 | 61 |
| Inflammation of central nervous system | 1.33E-19 | Increased | 2.333 | 102 |
| Invasive tumor | 1.09E-19 | Increased | 3.538 | 224 |
| Migration of lymphatic system cells | 8.48E-20 | Increased | 4.544 | 95 |
| Lymphocyte migration | 5.97E-20 | Increased | 4.62 | 94 |
| Organization of cytoskeleton | 5.36E-20 | Increased | 4.437 | 249 |
| Chemotaxis | 4.82E-20 | Increased | 4.815 | 123 |
| Migration of mononuclear leukocytes | 3.71E-20 | Increased | 4.931 | 99 |
| Proliferation of lymphocytes | 3.65E-20 | Increased | 3.292 | 150 |
| Proliferation of immune cells | 3.58E-20 | Increased | 3.303 | 158 |
| Proliferation of lymphatic system cells | 3.30E-20 | Increased | 3.653 | 159 |
| Homing of cells | 2.85E-20 | Increased | 4.933 | 128 |
| Interaction of phagocytes | 2.61E-20 | Increased | 3.318 | 64 |
| Quantity of lymphatic system cells | 1.12E-20 | Increased | 4.27 | 161 |
| Proliferation of blood cells | 8.94E-21 | Increased | 2.84 | 172 |
| Migration of phagocytes | 7.21E-21 | Increased | 5.059 | 82 |
| Proliferation of mononuclear leukocytes | 3.68E-21 | Increased | 3.384 | 154 |
| Quantity of lymphocytes | 1.41E-21 | Increased | 4.164 | 155 |
| Cell movement of mononuclear leukocytes | 7.82E-22 | Increased | 4.793 | 119 |
| Quantity of lymphoid cells | 7.71E-22 | Increased | 4.267 | 156 |
| Cell movement of myeloid cells | 9.41E-23 | Increased | 5.148 | 136 |
| Atherosclerosis | 5.83E-23 | Increased | 2.673 | 111 |
| Arteriosclerosis | 5.55E-23 | Increased | 2.673 | 112 |
| Cancer of cells | 5.08E-23 | Increased | 2.324 | 684 |
| Occlusion of artery | 8.44E-24 | Increased | 2.291 | 124 |
| Activation of leukocytes | 7.41E-24 | Increased | 3.906 | 152 |
| Quantity of mononuclear leukocytes | 3.21E-24 | Increased | 4.23 | 166 |
| Development of digestive organ tumor | 2.85E-24 | Increased | 2.223 | 805 |
| Occlusion of blood vessel | 1.78E-24 | Increased | 2.439 | 127 |
| Cell movement of phagocytes | 2.32E-25 | Increased | 5.16 | 143 |
| Vaso-occlusion | 1.80E-25 | Increased | 2.762 | 130 |
| Inflammatory response | 5.12E-26 | Increased | 4.721 | 179 |
| Neoplasia of cells | 4.32E-26 | Increased | 3.343 | 764 |
| Nervous system neoplasm | 1.09E-26 | Increased | 2.631 | 863 |
| Binding of leukocytes | 6.17E-27 | Increased | 4.377 | 107 |
| Adhesion of immune cells | 3.52E-27 | Increased | 4.659 | 102 |
| Activation of blood cells | 2.67E-27 | Increased | 4.18 | 171 |
| Binding of blood cells | 1.99E-28 | Increased | 4.151 | 117 |
| Activation of cells | 1.77E-28 | Increased | 4.235 | 212 |
| Adhesion of blood cells | 1.50E-29 | Increased | 4.576 | 110 |
| Cell movement of leukocytes | 2.69E-31 | Increased | 5.68 | 193 |
| Leukocyte migration | 2.22E-31 | Increased | 6.163 | 217 |
| Quantity of cells | 2.80E-32 | Increased | 4.371 | 346 |
| Migration of cells | 2.28E-33 | Increased | 5.915 | 387 |
| Quantity of leukocytes | 8.59E-34 | Increased | 3.627 | 215 |
| Quantity of blood cells | 1.14E-34 | Increased | 3.879 | 234 |
| Cell movement | 1.12E-34 | Increased | 6.237 | 422 |
| Digestive organ tumor | 2.14E-45 | Increased | 2.245 | 1243 |
| Intraabdominal organ tumor | 1.16E-50 | Increased | 2.485 | 1285 |
| Cancer | 3.98E-60 | Increased | 3.984 | 1378 |
| Solid tumor | 1.91E-61 | Increased | 2.377 | 1380 |
| Malignant solid tumor | 6.41E-62 | Increased | 2.227 | 1377 |
| Non-melanoma solid tumor | 2.74E-65 | Increased | 2.163 | 1366 |

**Supplemental Figure 1:**
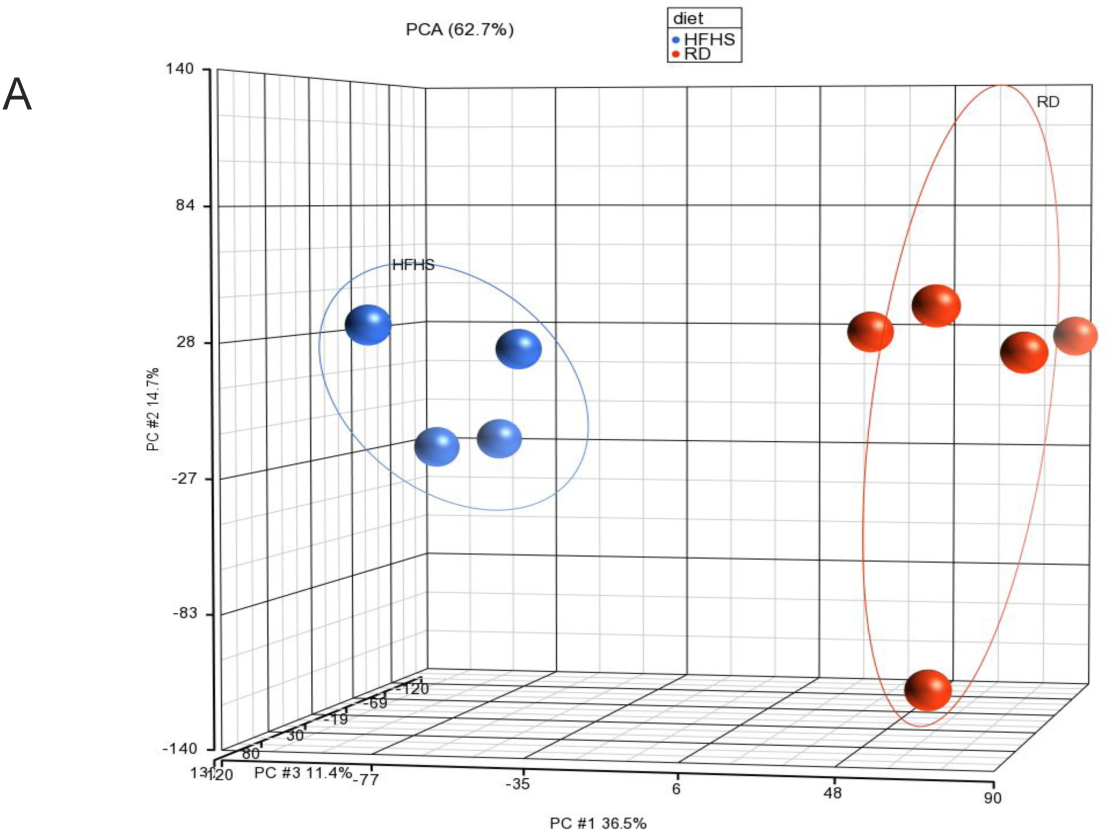
**A.** RNA was isolated for gene expression analyses from liver tissue at 16 weeks and principal component analysis performed. Colors refer to legend on top. Abbreviations: RD = regular diet, HFHS = high-fat high-sugar, PC = principal component.

**Supplemental Figure 2:**
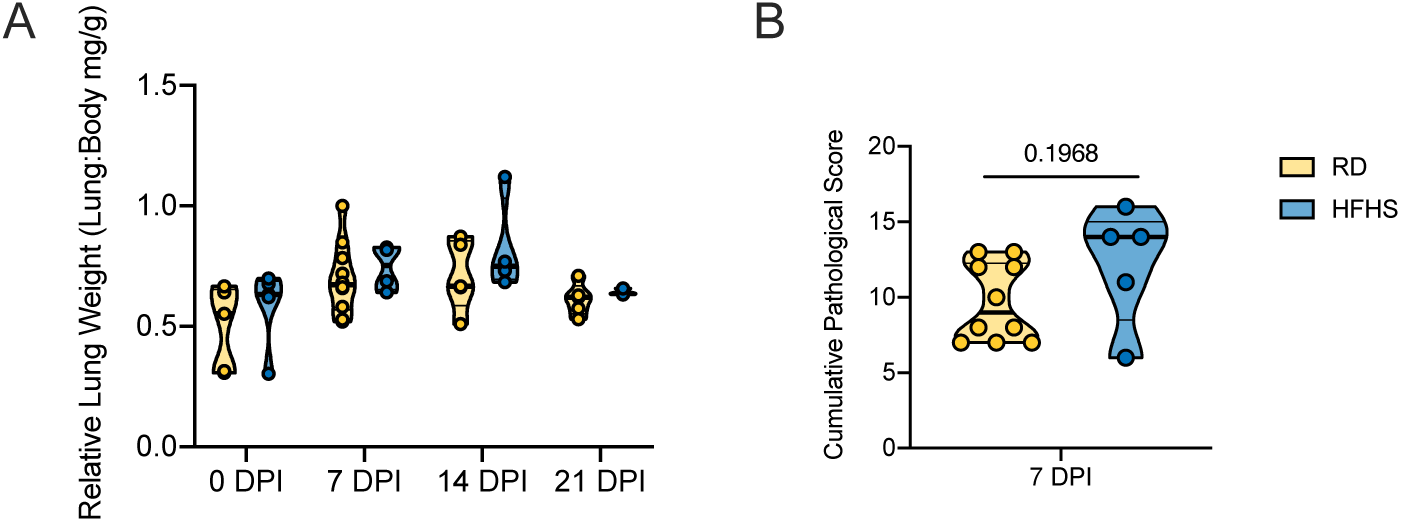
Male Syrian hamsters were fed either a regular or high-fat high-sugar *diet ad libitum* for 16 weeks, then challenged with 8×10^4^ TCID_50_ SARS-CoV-2. Animals were euthanized pre-challenge (0 DPI), at 7, 14 and 21 DPI. **A.** Lung weights. Truncated violin plots depicting median, quartiles, and individuals. **B.** Cumulative pathology score of lung tissues collected at 7 DPI. Truncated violin plots depicting median, quartiles and individuals, N = 10 (RD) / 4 (HFHS), Mann-Whitney test. Abbreviations: RD = regular diet, HFHS = high-fat high-sugar. p-values are indicated were appropriate.

**Supplemental Figure 3:**
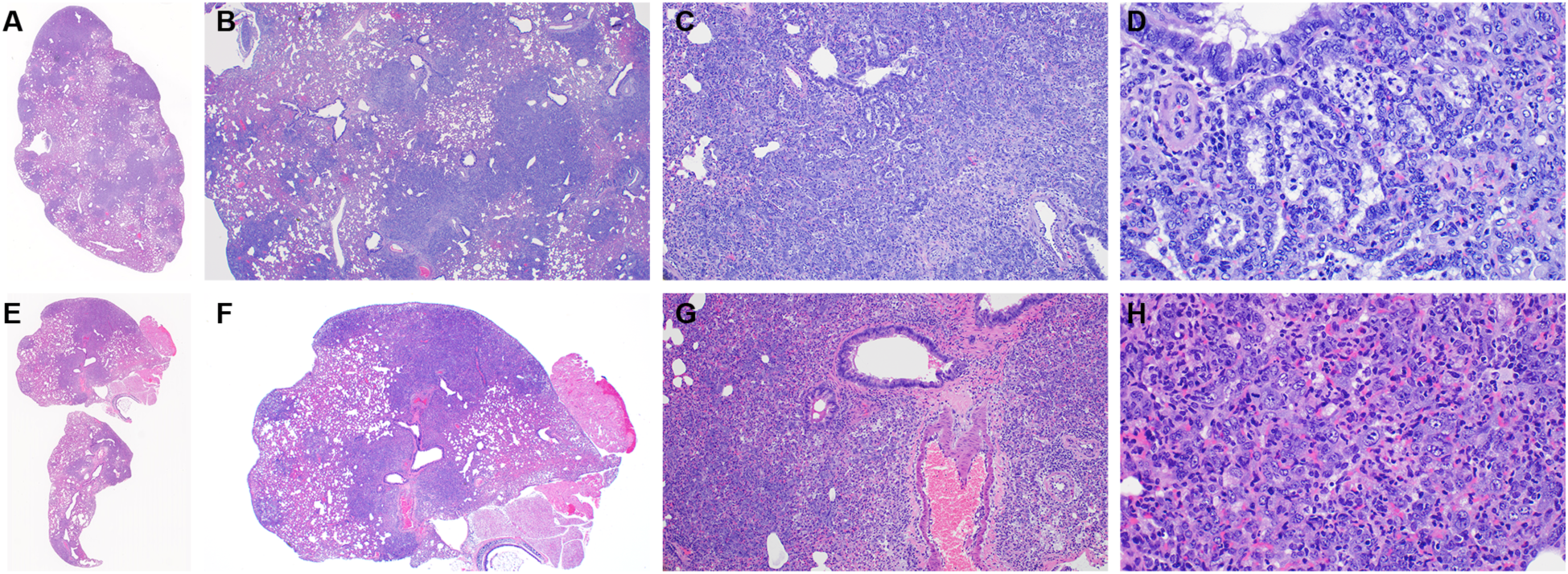
Male Syrian hamsters were fed either a regular or high-fat high-sugar *diet ad libitum* for 16 weeks, then challenged with 8×10^4^ TCID_50_ SARS-CoV-2. Animals were euthanized at day 8 and 9 due to increased weight loss. **A, E.** Dark, discreet foci identify areas of pneumonia; lighter areas indicate hemorrhage, edema, inflammation. HE, 1.4x. **B, F.** Although approximately 100% of the lobe is affected, only 50% contains discreet foci of interstitial pneumonia, HE, 20x. **C, D.** Examples of organized type II pneumocyte hyperplasia giving a honeycomb appearance. HE, 100x, 400x. **G, H.** Less well organized foci with more congestion, edema, and inflammation. HE, 100x, 400x. Of note, both appearances overlap and can be present in the same animal.

**Supplemental Figure 4:**
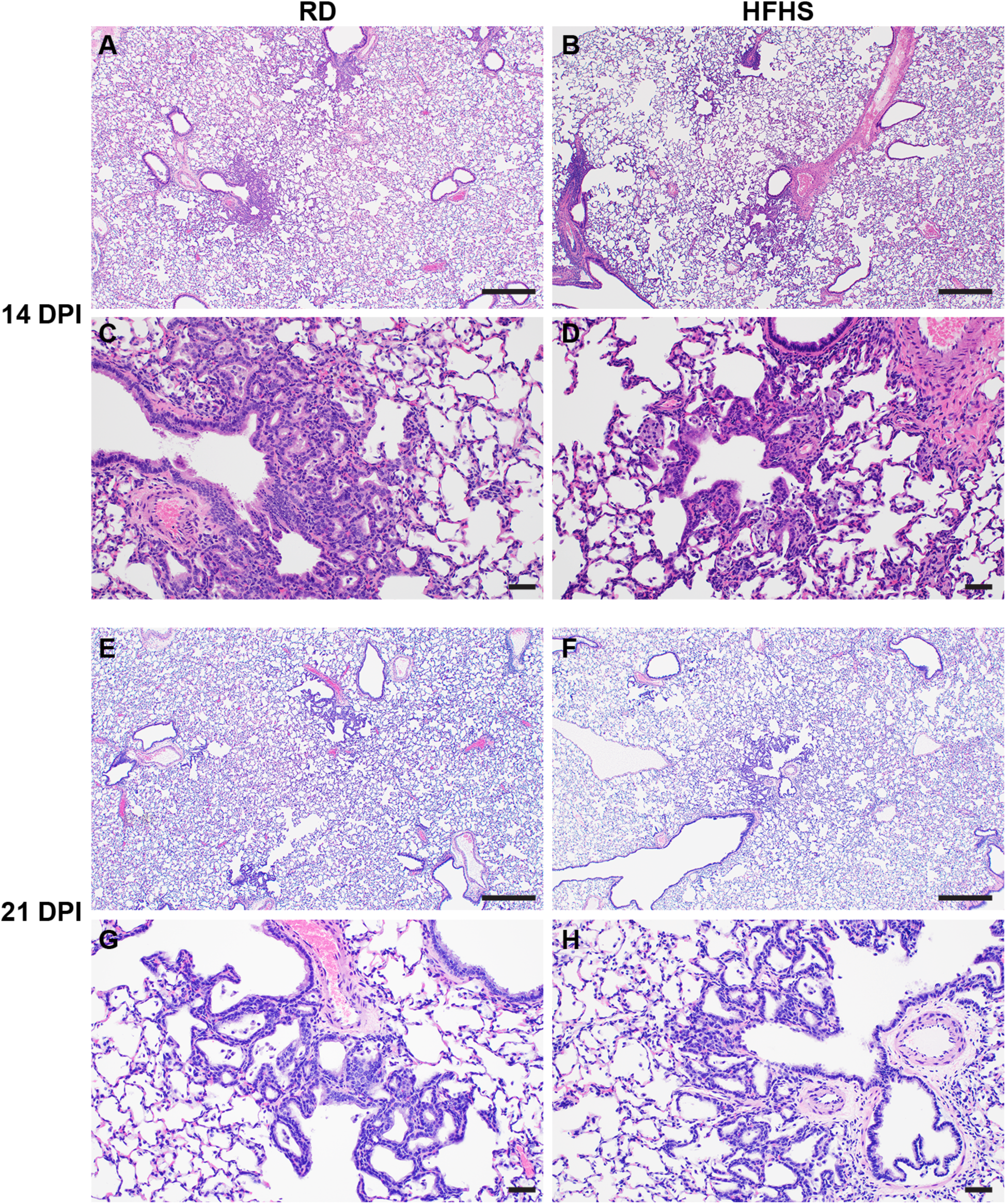
Male Syrian hamsters were fed either a regular or high-fat high-sugar *diet ad libitum* for 16 weeks, then challenged with 8×10^4^ TCID_50_ SARS-CoV-2. Lung tissues were collected 14 and 21 days post inoculation. **A, B**. 14 DPI, Lesions located at terminal bronchioles. HE, 40x. **C, D**. 14 DPI, Thickened septa, alveolar bronchiolization and minimal inflammation. HE, 400x. **E, F**. 21 DPI, Lesions appear indistinguishable. HE, 40x. **G, H**. 21 DPI, Thickened septa and alveolar bronchiolization remain. HE, 400x. Abbreviations: Reg = regular diet, HFHS = high-fat high-sugar, DPI = days post inoculation.

**Supplemental Figure 5:**
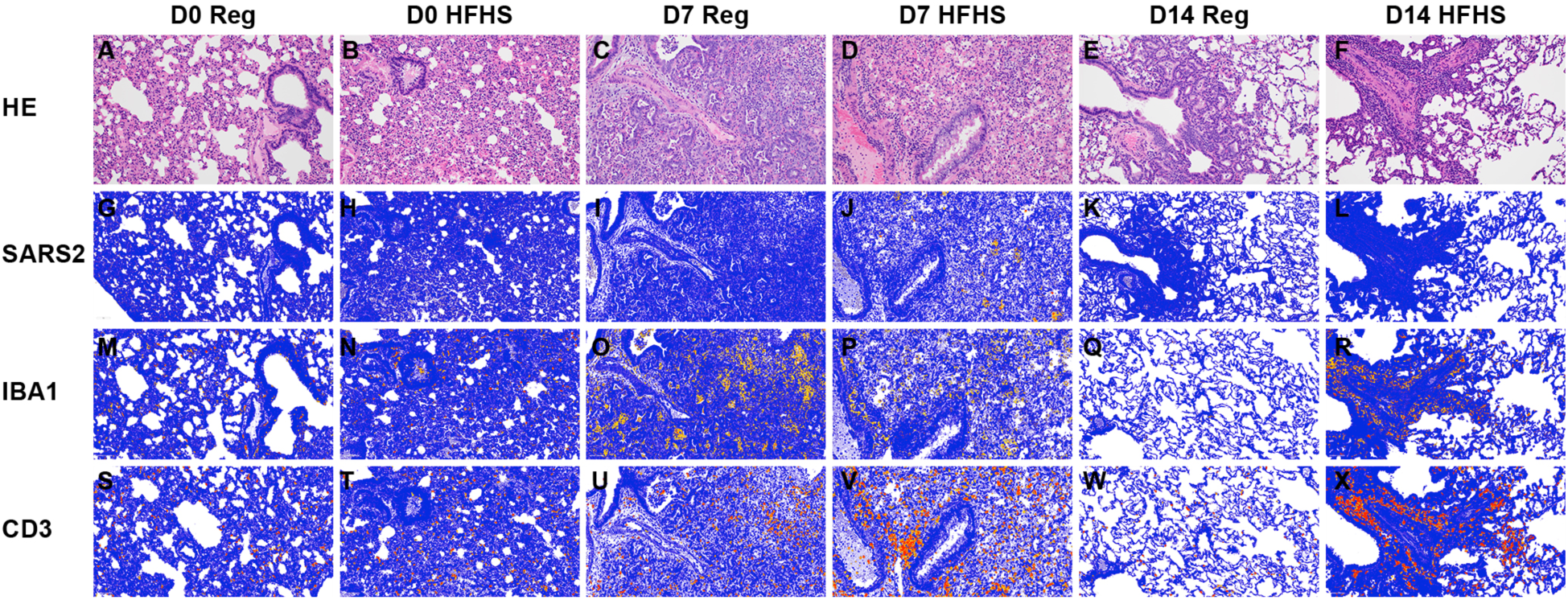
Male Syrian hamsters were fed either a regular or high-fat high-sugar *diet ad libitum* for 16 weeks, then challenged with 8×10^4^ TCID_50_ SARS-CoV-2. Animals were euthanized pre-challenge (0 DPI), 7 and 14 days post inoculation. Serial images of lungs. **A-F**. Pre-challenge lungs appear normal, 7 DPI lungs are pneumonic, and 14 DPI lungs appear to be resolving. HE, 200x. **G-L.** Positive pixel image of IHC staining against N protein of SARS-CoV-2. Note the positive pixels at 7 DPI in the HFHS image, 200x. **M-P.** Positive pixel image of IHC staining against IBA1. Note the increase in positive pixels at 7 and 14 DPI for both the RD and HFHS samples, 200x. **Q-X.** Positive pixel image of IHC staining against CD3, Note the increase in positive pixels at 7 and 14 DPI for both the RD and HFHS samples, 200x. Positive pixel = orange. Abbreviations: Reg = regular diet, HFHS = high-fat high-sugar, DPI = days post inoculation.

